# Fertilization-induced synergid cell death by RALF12-triggered ROS production and ethylene signaling

**DOI:** 10.1101/2024.03.10.584218

**Authors:** Junyi Chen, Huan Wang, Jinlin Wang, Xixi Zheng, Wantong Qu, Huijian Fang, Shuang Wang, Le He, Shuang Hao, Thomas Dresselhaus

## Abstract

Fertilization-dependent elimination of the persistent synergid cell is essential to block supernumerary pollen tubes and thus to avoid polyspermy in flowering plants. Little is known about the molecular mechanisms ensuring timely induction and execution of synergid cell death. We analysed manually isolated maize synergid cells along their degeneration and show that they are gland cells expressing batteries of genes encoding small secreted proteins under control of the MYB98 transcription factor. This network is down-regulated after fertilization, while genes involved in reactive oxygen species (ROS) production, ethylene biosynthesis and response, senescence, and oxidative stress regulation are induced before synergid elimination and its ultimate fusion with the endosperm. We further show that fertilization-induced RALF12 peptide specifically triggers mitochondrial ROS and apoptosis, while ethylene promotes synergid degeneration. In conclusion, this study sheds light on developmental programmed cell death (dPCD) in plants and provides a unique resource to discover novel PCD regulators.

## Introduction

Programmed cell death (PCD) is a widespread phenomenon that is essential for all multicellular organisms. It occurs in a highly controlled manner to selectively remove damaged or no longer needed cells that are sacrificed for the good of the whole organism^1–3^. PCD in plants is generally classified into environmental PCD (ePCD), which is triggered by external biotic or abiotic factors, and developmental PCD (dPCD), which occurs during vegetative and reproductive development^3^. dPCD is particularly frequent in plant reproduction^4^ and thus is crucial for fertilization success and seed production. During sex determination in many flowering plant species, carpel primordia cells are eliminated in male flowers, while stamen abortion occurs in female flowers. During microsporogenesis and megasporogenesis, tapetum cells and three of four megaspores are degenerated, respectively. PCD further occurs during nucellus and endosperm development but appears to be especially important for fertilization. During self-incompatible pollen-pistil interactions, PCD prevents alien and self-pollen tubes from growing toward ovules. Upon compatible pollen tube reaching the female gametophyte, the pollen tube cell and two synergid cells undergo degeneration in a highly regulated two-step process^5–7^. Ca^2+^ spiking in synergid cells and ROS-induced Ca^2+^ channel activation in the pollen tube lead to simultaneous burst of the pollen tube and one of the two synergid cells (i.e., receptive synergid), resulting in sperm cell release for double fertilization^8–10^. The second synergid cell (i.e., persistent synergid) remains intact during this step and maintains the capacity to attract additional pollen tubes in the case of fertilization failure^7,11,12^. Notably, polyspermy leads to lethal genome imbalance and chromosome segregation defects during embryo development^13^. The attraction of supernumerary pollen tubes is usually avoided. Once fertilization is achieved, the persistent synergid is removed in a second step with some delay to block supernumerary pollen tubes and prevent polyspermy. Ultimately, the persistent synergid cell fuses with the large fertilized central cell and is thus eliminated^14^. It has been discussed that removal of the persistent synergid cell is not achieved by PCD^4^.

Despite its abundance and importance for plant life, the gene regulatory networks (GRNs) controlling the (i) induction, (ii) effector, and (iii) degradation phases of dPCD are still poorly understood. This is mainly attributed to the occurrence of dPCD in embedded cells, which are surrounded by many other cells and thus are not accessible. Additionally, the initiation of dPCD is difficult to predict. During double fertilization in flowering plants, timely degeneration of synergid cells is crucial for reproductive success^5,6,14–18^. It has been shown that MITOGEN-ACTIVATED PROTEIN KINASE 4 (MPK4) prevents premature synergid cell death and FERONIA receptor kinase (FER)-controlled ROS accumulation^6,10^. A ‘cell death module’ consisting of REM transcription factors, VALKYRIE (VAL) and VERDANDI (VDD), both targets of the ovule identity MADS-box complex SEEDSTICK-SEPALLATA3, controls the death of the receptive synergid cell^19^. The plant hormone ethylene has been implicated in persistent synergid removal as the persistent synergid maintains its integrity longer in the ethylene-insensitive mutant *ein3 eil1*^17^. However, mutants incapable of producing ethylene show no PCD-related phenotype in synergid cells^18^ indicating that the ethylene signaling pathway, but not the hormone itself, might be involved in synergid cell death in *Arabidopsis*. It was suggested that an unknown ethylene-independent signaling pathway may control the death of the persistent synergid cell individually or synergistically with the ethylene-signaling pathway^18^. Nevertheless, the timely activation and execution of synergid degeneration and thus a fundamental principle in ensuring reproductive success remained unclear.

In this work, by overcoming technical limitations, we developed methods to isolate maize synergid cells at defined stages, then systematically explored the characteristic degeneration processes and the molecular machinery responsible for driving this system. We determined the exact timing of the loss of synergid-specific GRN and the induction of synergid PCD. PCD inducers, transcription factors, executors, as well as autophagy activities are highly activated a few hours before synergid-endosperm fusion (SE fusion). Moreover, we showed that the fertilization-induced autocrine signal ligand RALF12 specifically triggers mitochondrial ROS maximum at the filiform apparatus region to induce dPCD. This apoptosis pathway works synergistically with activated ethylene production and signaling in sensitized synergid cells, ensuring timely dPCD execution and corpse clearance shortly before SE fusion. In conclusion, this study uncovers a fundamental mechanism of dPCD during the essential fertilization process and provides a valuable resource for further studies investigating key molecular players responsible for driving dPCD in plants.

## Results and Discussion

### Timing of persistent synergid degeneration and generation of stage-specific RNA-seq data

To investigate the degeneration of the persistent synergid cell after fertilization in maize, we first monitored the morphological changes of synergid cells using a synergid cell marker line *pZmES4::ZmES4-GFP* (Fig. 1a–d)^20^ and additionally analyzed the inbred line B73 (Supplementary Fig. 1a–d). Before pollination, two synergid cells with comparable morphology were observed at the micropylar region of the female gametophyte (Fig. 1a and Supplementary Fig. 1a). 12 h after pollination (HAP), a dark and degenerated receptive synergid cell was visible, indicating that fertilization occurred (Supplementary Fig. 1b). The second synergid cell, termed the persistent synergid cell, was visible until 24-26 HAP (Fig. 1b, c). At about 25–28 HAP, the persistent synergid cell became abruptly invisible, leaving the degenerated receptive synergid and the zygote tightly attached at the micropylar region of the embryo sac (Supplementary Fig. 1c, d). Similarly, the synergid marker became no longer detectable at 28 HAP (Fig. 1d). These observations indicate that the persistent synergid cell has been completely removed at around 25–26 HAP, which corresponds to 17–18 h after fertilization (HAF), taking into consideration that it takes ∼ 8 h for pollen tubes to germinate and grow towards the female gametophyte at the silk length of about 10 cm^21^ (Fig. 1e–h).

**Fig. 1.**
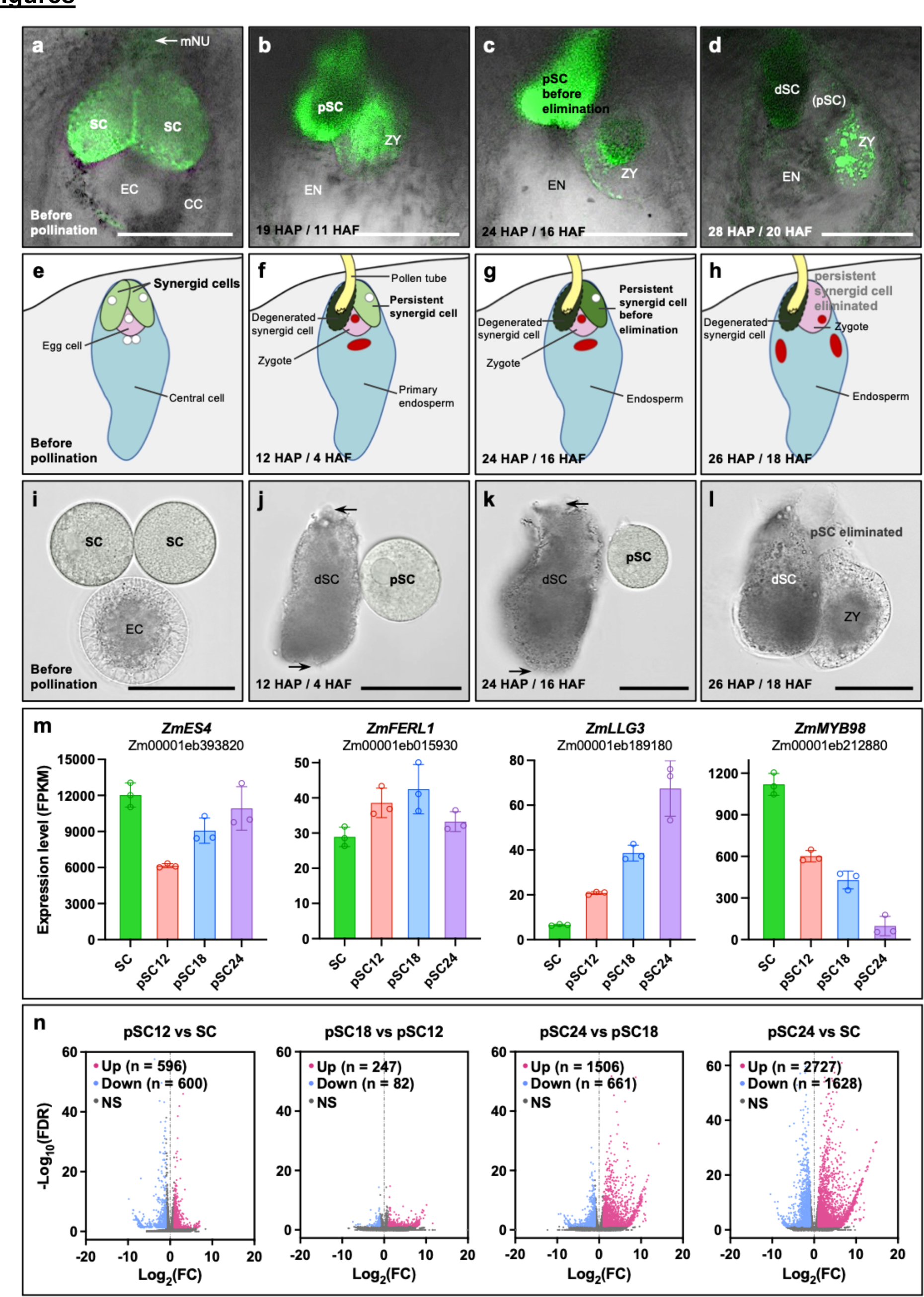
High-precision transcriptome changes during synergid degeneration in maize. **a**–**d** Tracing synergid cell degeneration using *pZmES4::ZmES4-GFP* as a marker line at indicated times before and after pollination/fertilization. *ZmES4* is expressed in synergid cells and weakly in the egg cell and zygote, respectively. At 28 HAP the synergid signal disappeared. **e**–**h** Scheme showing the timing of the double fertilization process and elimination of the two synergid cells. **i**– **l** Manually isolated synergid cells at indicated stages. At 26 HAP the persistent synergid cell is no longer visible and the zygote is tightly attached to the degenerated receptive synergid cell (l). Black arrows indicate pollen tube entrance and sperm cell release regions of the receptive synergid cell, respectively. **m** Expression pattern of known synergid-expressed genes at indicated synergid stages. Transcript levels are shown as FPKM (Fragments Per Kilobase of exon model per Million mapped fragments) values (means ± SD) of three biological replicates. **n** Volcano plots depict transcriptional dynamics of differentially expressed genes (DEGs) between indicated stages. Abbreviations: CC, central cell; dSC, degenerated synergid cell; EC, egg cell; EN, endosperm; HAF, hours after fertilization; HAP, hours after pollination; mNU, micropylar nucellus; pSC, persistent synergid cell; SC, Synergid cell; ZY, zygote. Scale bars, 50 μm.

Next, we developed methods to isolate living synergid cells and collect cells at defined stages after pollination/fertilization. Since many vacuoles were present in the peripheral region of the egg cells, they could be easily distinguished from synergid cells (Fig. 1i). Morphological changes of receptive synergid cells (they appeared dark after receiving pollen tube contents) facilitated discrimination of synergid cells before fertilization and persistent synergid cells shortly after fertilization (Fig. 1j and Supplementary Fig. 1b). Thereafter, synergid cells before pollination (SC), persistent synergid cells shortly after fertilization (pSC12, 12 HAP/about 4 HAF), during cell degeneration (pSC18, 18 HAP/ about 10 HAF) and shortly before cell elimination (pSC24, 24 HAP/about 16 HAF, i.e., only ∼ 1–2 h before cell deletion) were manually isolated (see Fig. 1i–l for isolated cells) and collected to generate high-precision degenerating stage-specific transcriptomic data at cellular resolution. Three independent biological replicates were prepared for each cell stage and sequenced at a depth of over 53 million reads per library (Supplementary Data 1). Comparison of the three biological replicates showed that the expression values between them were highly correlated (average *R*^2^ = 0.93). Principal component analysis (PCA) further separated these synergid cell samples into four groups (Supplementary Fig. 2). In total, transcripts of 14,700–15,800 unique genes were detected in synergid cells at sequential developmental stages (Supplementary Data 1 and 2).

For initial analysis of dynamic gene expression during the synergid cell degeneration processes, we focused on known synergid-expressed genes (Fig. 1m). *ZmES4*, encoding secreted peptides required for pollen tube burst and sperm cell release, was highly expressed in synergid cells and significantly downregulated shortly after fertilization as described previously^20^. The *Arabidopsis* receptor-like protein kinase FERONIA and its coreceptor LORELEI were shown to act specifically at the synergid cell surface for pollen tube reception^22,23^. Their orthologs in maize, *ZmFERL1* and *ZmLLG3*^24^, were expressed in synergid cells and showed distinct activation patterns during synergid cell degeneration. We also found that *ZmMYB98*, the most highly expressed transcription factor gene in maize synergid cells, exhibited a substantial and continuous decrease during synergid degeneration (Fig. 1m and Supplementary Data 5). In summary, the observed gene expression patterns in synergid cells are in line with previous reports and expectations, which together with strong correlation between independent biological replicates assure the high quality and reliability of the data.

To gain further insight into the timing and scale of PCD triggering, activation, and execution in the persistent synergid cell, we identified significant upregulation of 596 genes in pSC12, 247 genes in pSC18, and 1506 genes in pSC24 (Fig. 1n, Supplementary Fig. 3 and Supplementary Data 7). In total, we observed significant upregulation of 2,727 genes in persistent synergid cells at 24 HAP compared to synergid cells. Substantial changes were observed between the transcriptional profiles of pSC18 and pSC24, indicating that the major PCD-related activation wave occurs later after fertilization and only within a few hours before persistent synergid elimination.

### Cell-specific gene expression in synergid and egg cells

Next, we analyzed gene expression patterns in synergid cells of maize. Among the top 30 genes with highest expression values, 26 genes (86.7%) encode proteins with predicted N-terminal signal peptides (SPs), indicating that they are targeted to the secretory pathway. In contrast, in the neighboring egg cells/zygotes, only 10% of the top 30 genes encode proteins carrying SPs for protein secretion (Fig. 2a and Supplementary Data 3). Remarkably, in the TOP200 category, substantially more synergid cell genes encode ER/Golgi localized proteins (11.5% in synergid cells versus 1.5% in egg cells; Fig. 2b and Supplementary Data 3). Conversely, egg cells display remarkably enriched transcripts for mitochondrial proteins (Fig. 2b), indicating elevated mitochondrial biogenesis and/or activity in the female gamete. This result is consistent with the concept that maternal mitochondrial function is important for egg cell maturation and embryogenesis initiation^25^ in flowering plants. Besides, egg cells and zygotes contain comparable highly abundant ribosomal transcripts, which are orders of magnitude higher compared with synergid cells (Fig. 2b). This is consistent with previous studies in animal species showing that large amounts of ribosomes/ribosomal transcripts are produced and stored in the female gamete before fertilization, to fulfill efficient protein production during embryogenesis initiation. Taken together, these results show that synergid cells possess typical characteristics of glandular cells for protein secretion, while neighboring egg cells are well prepared for embryonic initiation.

**Fig. 2.**
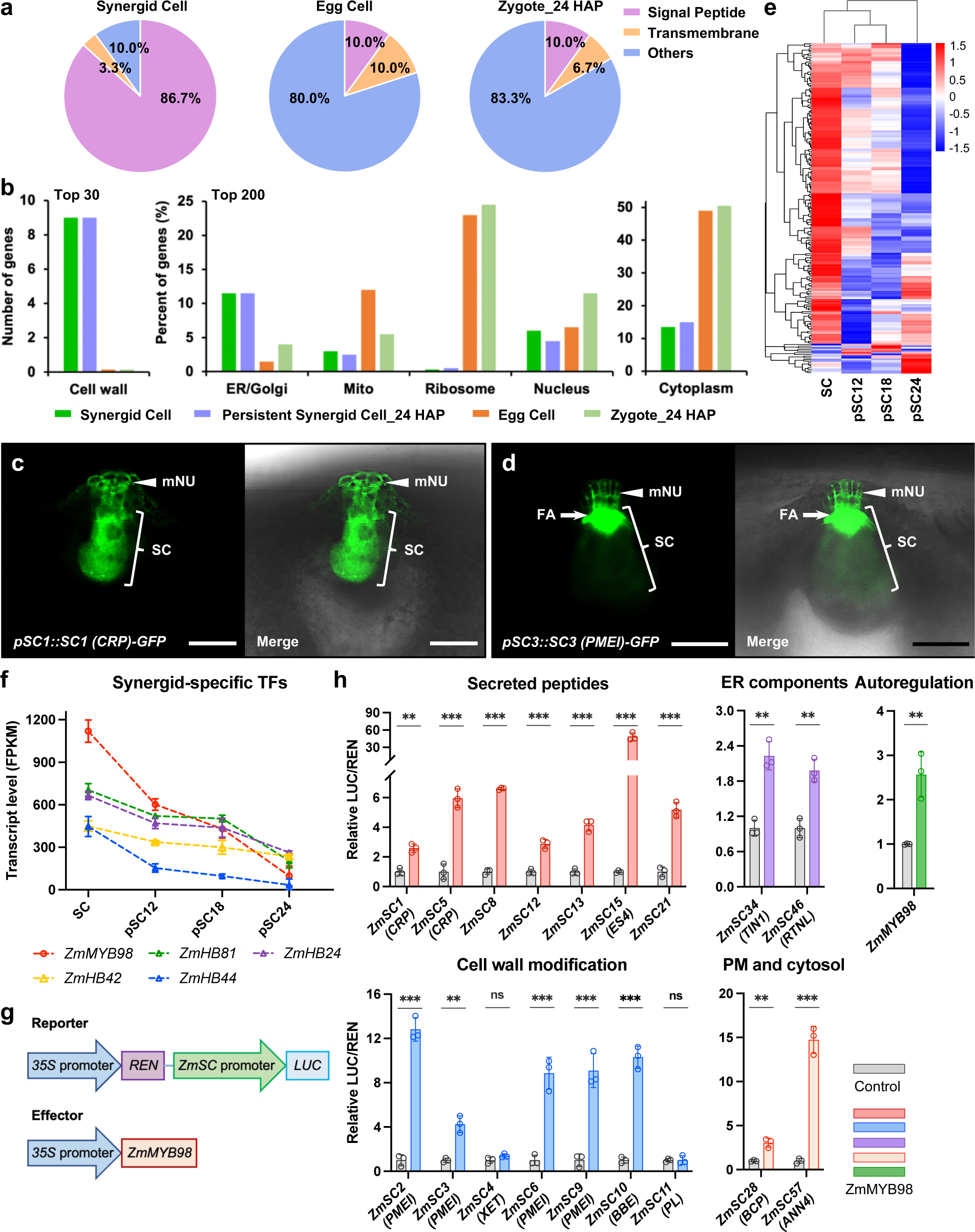
Synergids are glandular cells generating numerous peptides via a ZmMYB98 controlled GRN. **a** Proportion comparison of the 30 most highly expressed genes in synergid cells. 86.7% genes encode proteins containing predicted signal sequences for targeting to the secretory pathway. Egg cells and zygotes at 24 HAP are included for comparison. **b** Subcellular distribution of most abundant gene products in synergid cells, persistent synergid cells (24 HAP), egg cells, and zygotes (24 HAP). **c**–**d** Localization of the fusion proteins SC1-GFP (a CRP) (c) and SC3-GFP (a PMEI) (d) in synergid cells and cell walls of micropylar nucellus cells in mature embryo sacs before pollination. Arrowheads point to secreted signals at the micropylar nucellus region. **e** Heatmap showing expression pattern of the TOP200 synergid cell-expressed genes during the progression of synergid degeneration. **f** Expression analysis of the most abundant synergid cell-expressed transcription factors during synergid degeneration. **g** Scheme of reporter and effector vectors for dual-luciferase transactivation assays shown in (h). **h** Transactivation assay showing that ZmMYB98 positively regulates a comprehensive set of most highly expressed synergid-specific genes. Relative ratio of firefly luciferase (LUC) to *Renilla* luciferase (REN) was used to determine relative promoter activation activity. Results are an average of three independent experiments. Statistical significance is determined by Student’s *t*-test (\*\**P* < 0.01, \*\*\**P* < 0.001) and the error bars show standard deviation (SD). Abbreviations: FA, filiform apparatus; mNU, micropylar nucellus; SC, synergid cell. Scale bars, 50 μm.

Remarkably, 28 of the top 30 synergid cell genes (93.3%) show synergid-specific/predominant expression patterns (Supplementary Data 4). This high percentage of cell-specific expression of the most abundant transcripts was not observed in egg cells or zygotes, supporting the notion of highly specialized functions of flowering plant synergid cells. Moreover, the top synergid cell-specific genes exhibit extraordinarily high transcript levels, with 21 of them showing FPKM (Fragments Per Kilobase of exon model per Million mapped fragments) values greater than 10,000. We found that among the top 30 synergid cell genes, nearly half (14 out of 30 genes) encode small secreted peptides (Supplementary Data 4), including ZmESs with functions in pollen tube rupture during pollen tube reception^20,26^, and other synergid-specific cysteine-rich peptides (CRPs) with suggested functions in cell-cell recognition during double fertilization^27^. Notably, one-third (10 out of 30 genes) encode secreted cell wall modifiers, including four pectin methylesterase inhibitors (PMEIs) and three pectate lyases (PLs). We selected two previously undescribed genes (one encoding a novel CRP and one encoding a PMEI) for validation by expressing *pSC::SC-GFP* constructs in transgenic lines. Being consistent with the cell-specific gene expression data, both of them were strongly and specifically expressed in synergid cells, and were secreted to the micropylar region of the mature ovule (Fig. 2c, d). Further characterization of these abundantly and predominantly expressed secreted proteins will find out whether they are involved in facilitating double fertilization and ensuring reproductive success in the monocot and crop plant maize.

### Loss of synergid-specific GRN during synergid degeneration

In accordance with the observation that most highly expressed genes in synergid cells showed synergid-specific expression patterns (Supplementary Data 4), we found that eight of the top 10 synergid-expressed TFs were also specifically or predominantly expressed in synergid cells (Supplementary Data 5). Notably, their expression levels were significantly downregulated after fertilization during synergid degeneration (Fig. 2f and Supplementary Data 5). In particular, the expression of *ZmMYB98* was substantially decreased from a 1,200 FPKM value in synergid cells to 98 in persistent synergid cells shortly before cell deletion. A similar pattern was also found for the majority of the top 200 synergid-expressed genes (Fig. 2e and Supplementary Data 6).

Based on the consistent overall expression trend, we next asked whether there is a positive regulation between the top synergid-specific TFs and the top synergid-specific genes. As *ZmMYB98* was identified as the most strongly expressed TF gene in maize synergid cells, we hypothesized that it could regulate other synergid-specific genes (including itself). Therefore, we established a dual-luciferase assay to test whether ZmMYB98 can activate a selected group of most highly expressed synergid-specific genes (Fig. 2g). As shown in Fig. 2h, ZmMYB98 stimulated the activities of nearly all selected gene promoters except that of two genes involved in cell wall modification. All secreted peptide genes, including the most strongly expressed CRP genes with potential roles in pollen tube attraction^12,28^ and *ZmES4*^20^ were activated by ZmMYB98.

Moreover, four cognate genes encoding secreted cell wall modifier PMEIs, with proposed function in pollen tube growth arrest and reception^29^, exhibited strong induction by ZmMYB98 (Fig. 2h). In addition to secreted peptides and cell wall modifiers, two genes encoding synergid cell-predominant ER protein orthologs (ZmTIN1 and ZmRTNL), a gene encoding a synergid cell-specific plasma membrane blue copper protein (ZmBCP), and a gene encoding a synergid cell-specific cytoplasm annexin (ZmANN4), were also positively regulated by ZmMYB98. Since the *ZmMYB98* promoter region itself contains several putative MYB98 binding sites (TAAC)^30^, we then tested whether ZmMYB98 could regulate its own expression. The transactivation assay showed that *ZmMYB98* is indeed autoregulated (Fig. 2h and Supplementary Fig. 4).

To further test this regulation, we investigated the expression and subcellular localization of ZmMYB98-activated ZmSC-GFP fusion proteins under the control of their endogenous promoters in a heterologous dicot system *in planta* (Supplementary Fig. 4). In the absence of ZmMYB98, GFP signals could not be detected in transfected tobacco epidermal cells. In contrast, ZmMYB98 co-expression efficiently promoted the activity of synergid-specific promoters for ZmSC-GFP expression and secretion as indicated by GFP signals localized at the apoplastic space (Supplementary Fig. 4). These results confirm that the most highly expressed synergid-specific genes require ZmMYB98 for expression, and ectopic expression of ZmMYB98 in tobacco epidermal cells is sufficient to activate the expression of maize synergid-specific genes in these cells. We therefore conclude that ZmMYB98 controls the expression of most abundantly and predominantly expressed genes in synergid cells. This synergid-specific GRN is significantly downregulated during synergid degeneration and nearly shut off shortly before synergid cell elimination.

### Fertilization-induced ZmRALF12 triggers ROS production at the filiform apparatus region

ROS are signaling molecules for many basic biological processes, and moreover, they act as main triggers for physiological or programmed pathways for cell death during plant development, reproduction, and stress responses^31–33^. To determine whether ROS are involved in the degeneration process of the persisting synergid cell, we stained maize ovule tissues at different time points before and after fertilization, with the general ROS dye 2’,7’-dichlorofluorescein diacetate (DCFH-DA). Before pollination, ROS accumulation was detected in epidermal micropylar nucellus cells and was almost absent from mature synergid cells (Fig. 3a). However, at 20 HAP (∼12 HAF), we detected very intense DCFH-DC staining in the persisting synergid cell, with the maximum ROS accumulation adjacent to the plasma membrane (Fig. 3b–d, g). These findings indicate that fertilization induces ROS production accumulates to high levels before persistent synergid degeneration.

**Fig. 3.**
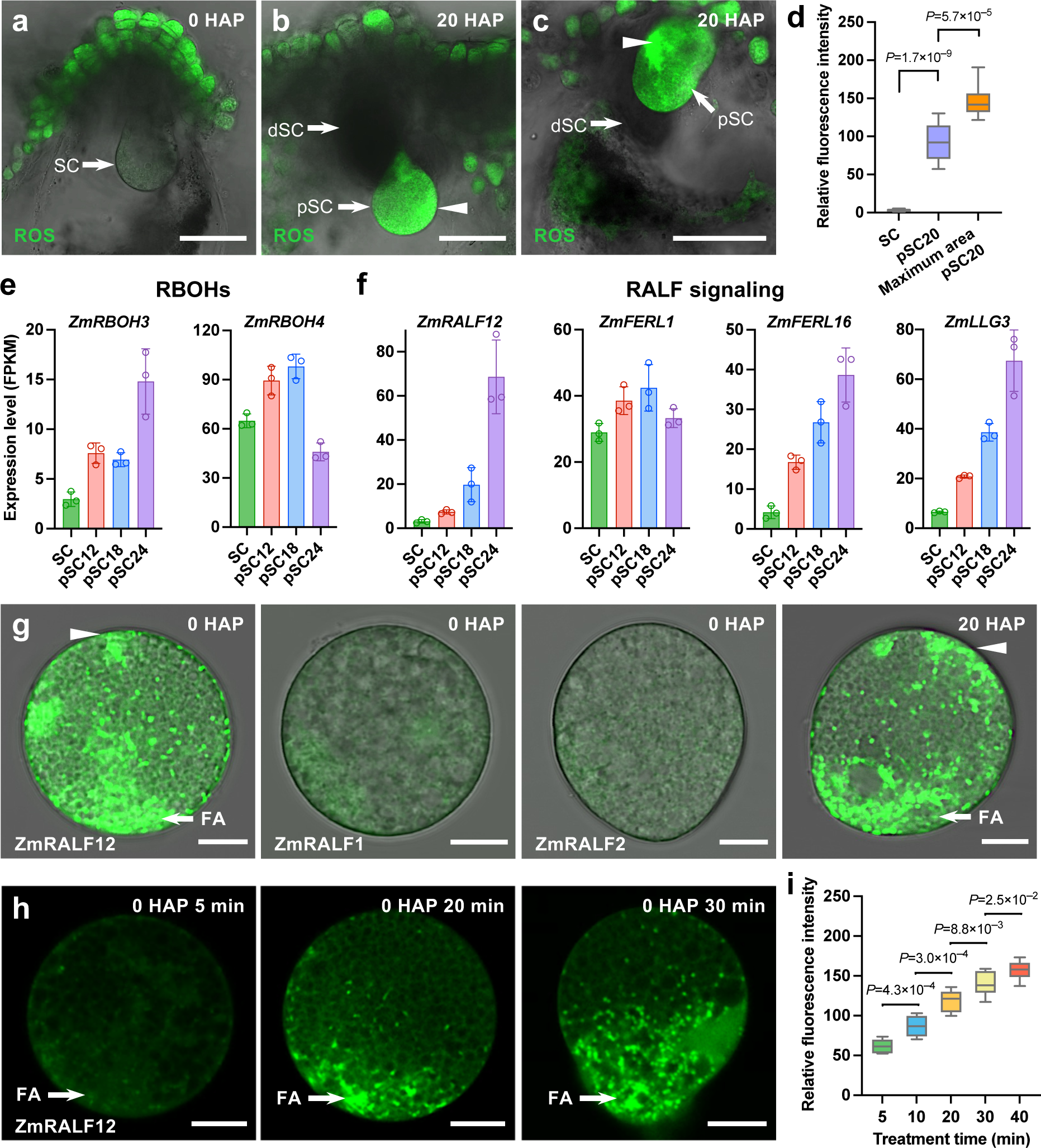
Fertilization-induced RALF12 triggers high levels of ROS production and accumulation at the filiform apparatus region of the persisting synergid cell. **a** DCFH-DA staining showing ROS accumulation in epidermal micropylar nucellus cells and its absence inside the mature embryo sac. **b**,**c** At 20 HAP strong staining is detectable inside the persistent synergid cell. Arrowheads point to ROS maxima in persistent synergid cells. **d** Quantification of ROS fluorescence intensity in synergid cells at stages shown in (a–c) (n=10). **e**,**f** Expression of RBOH and RALF peptide/receptor genes during fertilization and persistent synergid degeneration. **g** ZmRALF12, but not ZmRALF1 and ZmRALF2 induces ROS production and accumulation in synergid cells. After RALF12 application (4 μM for 30 min), synergid cells displayed strong and uneven ROS accumulation (i) comparable to persistent synergid cells at 20 HAP (j). **h** ZmRALF12 triggers rapid (within minutes) and specific (granular) ROS accumulation at the filiform apparatus (FA) region. **i** Quantification of ROS fluorescence intensity at the FA region after ZmRALF12 (4 μM) application at indicated time points (n=8). In box-and-whisker plots, centre lines represent the 50th percentile; bottom and top of each box indicate 25th and 75th percentiles, respectively; whiskers represent minimum and maximum, respectively. Statistical significance is determined by Student’s *t*-test. Abbreviations: dSC, degenerated synergid cell; FA, filiform apparatus; pSC, persistent synergid cell; SC, synergid cell. Scale bars: 50 μm (a–c) and 10 μm (g, h), respectively.

Plant NADPH oxidases, also known as respiratory burst oxidase homologs (RBOHs), have been called ‘the engines of ROS signaling’^33^. Two *RBOH* genes, *ZmRBOH3* and *ZmRBOH4,* were expressed in maize synergid cells and were further transcriptionally activated at 12 HAP (Fig. 3e), indicating that major components of the ROS production machinery were present. RBOHs were previously shown to be activated by Rapid Alkalinization Factor (RALF) signaling peptides involving also members of the *Catharanthus roseus* RLK1-like (CrRLK1L) receptor kinase family and their GPI-anchored LORELEI-like (LLG) co-receptors^34,35^. Among the 24 RALF genes in maize^24^, only *ZmRALF12* was highly induced during persistent synergid degeneration (Fig. 3f and Supplementary Data 8). Additionally, two CrRLK1L and one LLG encoding genes (*ZmFERL1*, *ZmFERL16* and *ZmLLG3*) that potentially form a heterotypic receptor complex with the mobile RALF ligand^24^ were activated shortly after fertilization and continuously increased (with the exception of *ZmFERL1*) indicating that a functional RALF-CrRLK1L-LLG complex can be formed before synergid cell degeneration. We hypothesized that the fertilization-induced ZmRALF12 peptide ligand is the limiting factor that triggers ROS production and subsequent oxidative burst in the persisting synergid cell. To test this hypothesis, we applied each 4 μM ZmRALF12 and two pollen tube-expressed ZmRALF1 and ZmRALF2 peptides^24^ for 30 min to synergid cells isolated from virgin ovules. Intracellular ROS accumulation was detected using DCFH-DC staining. In comparison to ZmRALF1 and ZmRALF2, ZmRALF12 (gene ID Zm00001eb410350) treated synergid cells displayed strong and uneven ROS accumulation comparable to persistent synergid cells at 20 HAP (Fig. 3g). We observed that ROS were prominently located at the filiform apparatus and at discrete aggregation near the periphery of the plasma membrane, with typical granular compartmentation likely representing peroxisomes and mitochondria, respectively, associated with ROS production, accumulation and signaling (Fig. 3g).

To further explore the temporal and spatial dynamics of ZmRALF12-induced ROS production, we generated a time series of ZmRALF12 treatment on synergid cells, ranging from 5 to 40 min. Remarkably, even at 5 and 10 min, respectively, ZmRALF12 triggered granular ROS accumulation in close proximity to the filiform apparatus membrane of the synergid cells (Fig. 3h and Supplementary Fig. 5a), demonstrating rapid RALF-induced ROS production. Extended periods of ZmRALF12 incubation led to significantly elevated granular ROS accumulation at this region and an ultimate granular ROS expansion throughout the whole synergid cell (Fig. 3h, i and Supplementary Fig. 5a). Similarly, increased ZmRALF12 concentrations correlated with higher ROS levels at the filiform apparatus area of the synergid cells and the same trend of granular ROS diffusion towards their opposite cellular domain (Supplementary Fig. 5b,c). Taken together, we conclude that fertilization-induced synergid cell-derived autocrine signal ZmRALF12 rapidly (within minutes) triggers granular ROS accumulation at the filiform apparatus region, and mediates high ROS levels in a time-and concentration-dependent manner.

### ZmRALF12 triggers mitochondrial ROS production, oxidative stress, and dPCD in the persisting synergid cell

To further investigate subcellular ROS production and accumulation during synergid cell death, we manually isolated persistent synergid cells at 20 HAP and double-stained the cells with a mitochondria-specific fluorescence dye (Mito-Tracker Red) and the ROS marker DCFH-DA. We found both, ROS and mitochondrial signals, showed prominent localization as accumulated particles near the filiform apparatus (Fig. 4a). Signals were densely populated at the cellular synergid domain that also contains the FER-LLG receptor complex in the model plant *Arabidopsis*^22,23^. Colocalization of ROS production sites and mitochondria was revealed by the overlap of signals resulting in yellow staining and a strong correlation of relative fluorescence intensity plots (Fig. 4b). The same result was obtained when double staining was performed on ZmRALF12-treated synergid cells (Fig. 4c, d). Notably, in both cases, the strongest ROS production area was at the filiform apparatus region (i.e., the cellular ZmRALF12-receptor interacting region). A decrease in ROS signal intensity correlates well with a longer distance from the micropylar filiform apparatus (Fig. 4b, d). Collectively, these results indicate that fertilization-induced ZmRALF12 triggers ROS production and accumulation in synergid mitochondria from the filiform domain of synergid cells.

**Fig. 4.**
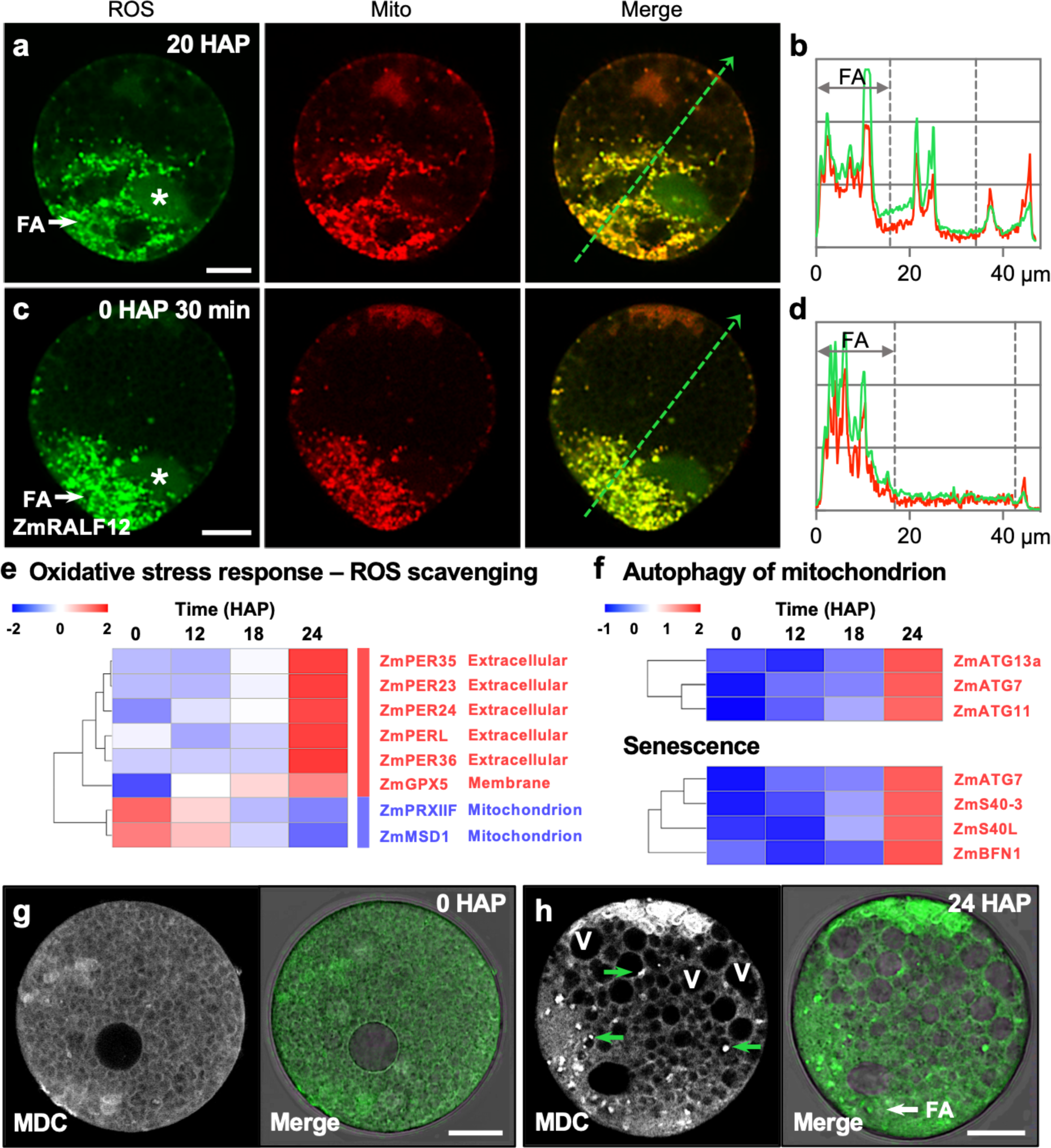
Fertilization-induced autocrine signal RALF12 mediates mitochondrial oxidative stress and synergid apoptosis. **a**,**b** High levels of ROS accumulation in mitochondria at the filiform apparatus region of the persisting synergid cell at 20 HAP. **c**,**d** RALF12 application (4 μM for 30 min) triggers mitochondrial ROS accumulation in synergid cells comparable to persistent synergid cells at 20 HAP. Asterisks indicate moderate ROS accumulation in synergid nuclei (a,b) and merged images of ROS accumulation and a mito-tracker are as indicated (a, c). Intensity plots (b, d) were generated along the green and interrupted arrows in (a, c). **e** In contrast to the activation of extracellular targeted ROS scavenging enzymes, mitochondrial ROS scavenging enzymes are downregulated after fertilization and continually decreased during synergid degeneration. **f** Mitophagy and senescence-related genes are strongly activated at 24 HAP. **g**,**h** Monodansylcadaverine (MDC) staining showing an intact synergid cell before fertilization (g) and active autophagic processes and vacuole accumulation in the persistent synergid cell before its elimination (h). Green arrows point to autophagosomes transported to vacuoles. White arrow shows autophagy maxima at the mitochondrial ROS accumulating filiform apparatus region. Scale bars, 10 μm.

Consistent with a strong increase of *ZmRALF12* expression in synergid cells from 18 to 24 HAP and the observed mitochondrial ROS accumulation at 20 HAP, a subset of genes involved in peroxisomal functions including NAD^+^ import (*ZmPXN*), 2-hydroxyacid oxidation (*ZmGLO3*), very long-chain fatty acids (VLCFAs) synthesis for peroxisomal metabolization (*ZmKCR1*) and fatty acid oxidation (*ZmSDP1*, *ZmECH2* and *ZmKAT5*), were strongly activated in the persisting synergid cell after 18 HAP (Supplementary Fig. 6). These findings further indicate that the onset of oxidative burst occurs with some delay after fertilization, which ultimately provokes a shift from the compartmental redox balance towards oxidative stress. This shift was reported to be largely mediated through Ca^2+^ signaling^36^. Such a switch was evidenced by the activation of genes encoding plasma membrane and ER-membrane Ca^2+^ channels (*ZmCSC1* and *ZmTMCO1*), mitochondrial inner membrane Ca^2+^ uniporter that mediates Ca^2+^ uptake into mitochondria (*ZmMCU2*), and Ca^2+^ signaling components involved in senescence regulation (Supplementary Fig. 7). The concomitant induction of gene batteries for oxidative stress response and senescence (Fig. 4e, f) further confirms the shift from a balanced to an oxidative stress status in persisting synergid cells at ∼24 HAP.

Strikingly, in contrast to the upregulation of oxidative stress response ROS-scavenging enzymes that are targeted to the extracellular matrix, mitochondrial antioxidant enzymes, which protect cells from apoptosis^37,38^, are specifically downregulated after fertilization and during the progressing of synergid degeneration (Fig. 4e). By using monodansylcadaverine (MDC) staining, we further observed active autophagic processes and vacuole accumulation shortly (i.e., only ∼1– 2 h) before persistent synergid elimination (Fig. 4g, h). Notably, autophagosomes are also most abundant in the filiform apparatus region, where ZmRALF12-triggered mitochondrial ROS production and oxidative stress/damage are most evident (Fig. 4h). This is consistent with the activation of a series of mitophagy-related genes known to be involved in autophagic removal of oxidatively damaged mitochondria^39^ (Fig. 4f and Supplementary Fig. 8). Collectively, these results show that the peptide ligand ZmRALF12 triggers mitochondrial ROS production after successful fertilization accompanied with downregulation of mitochondrial antioxidant enzymes. This mediates the switch to oxidative stress and senescence thereby activating the dPCD pathway in the persisting synergid cell.

### Ethylene promotes sensitized synergid cell death

ROS signaling is interconnected with the response to several phytohormones, especially the gaseous hormone ethylene. Production of ethylene and the activation of ethylene responses have been reported to belong in vegetative development as the first events following the accumulation of ROS^40,41^ In *Arabidopsis*, it has been previously shown that ethylene signaling^17,18,42^, bot not necessarily the hormone itself^18^, is required for the degeneration of the persistent synergid cell and the establishment of a pollen tube block. To dissect the contributions of ethylene and ethylene signaling to synergid cell death, we therefore first investigated the dynamic expression changes of genes related to ethylene biosynthesis and signaling. The ethylene biosynthetic pathway consists of two well-defined enzymatic steps: first, ACC synthase (ACS) catalyzes the conversion of *S*-adenosyl methionine (SAM) to the intermediate 1-aminocyclopropane-1-carboxylic acid (ACC), and second, ACC oxidase (ACO) oxidizes ACC to generate ethylene^43,44^. Our cell-and stage-specific RNA-seq data set revealed that both *ACS* and *ACO* genes were expressed throughout the processes of fertilization and persistent synergid degeneration, but in contrast to *ACSs*, *ACOs* (*ZmACO5* and *ZmACO20*) showed a significant increase at 24 HAP (Fig. 5a and Supplementary Data 8). Accordingly, the key regulators of ethylene signaling, EIN3 and EIL1, which are sufficient for the activation of the ethylene-response pathway^45^, were *de novo* expressed after fertilization and showed an enormous increase at this later stage (Fig. 5a). In consequence, eight potential downstream AP2/ERF transcription factors were specifically activated in a similar pattern (Fig. 5a). Notably, in neighboring zygotes at 24 HAP, all eight AP2/ERF transcription factor genes as well as those encoding ACOs remained not expressed or unactivated^21^ indicating that the synergid cell-specific GRN is inactive in the neighboring zygote.

**Fig. 5.**
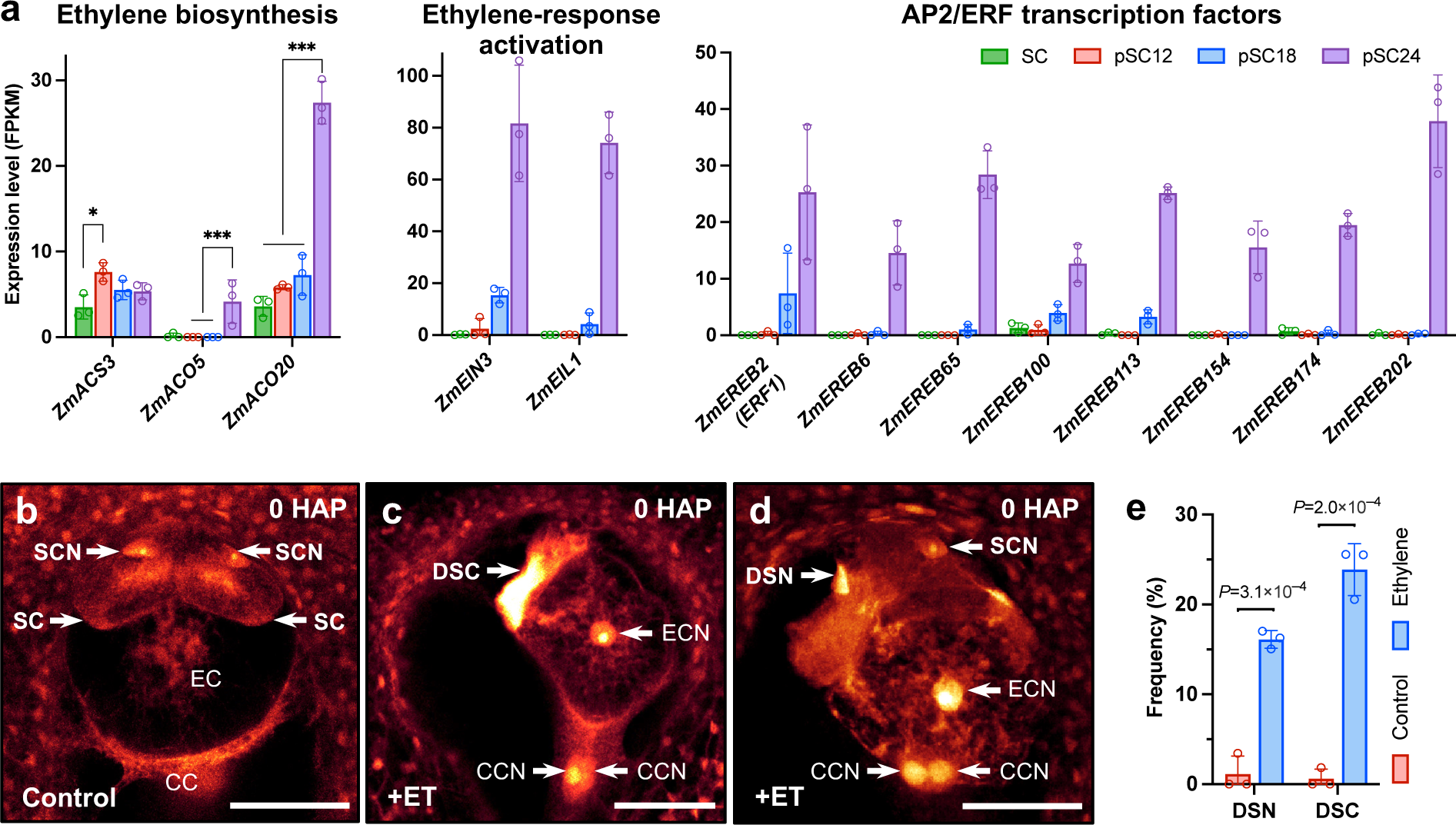
Post-fertilization ethylene production activates the ethylene signaling pathway and promotes synergid cell death in maize. **a** Strong activation of ethylene (ET) biosynthesis and ET-responsive genes at 24 HAP in the persistent synergid cell. For ET biosynthesis genes, significant activations are indicated (*Log_2_FC > 1 and adjusted *P* < 0.05, ***Log_2_FC > 1 and adjusted *P* < 0.001). **b**–**d** ET application specifically triggers only synergid cell death in the mature female gametophyte. CLSM of a female gametophyte without ET treatment (b; control) and 30 h after ET-treatment exhibiting fully degenerated synergid cells (c) and degenerated synergid nuclei (d), respectively. **e** Frequencies of synergid defects (control: n = 119; +ET: n = 163). Statistical significance is determined by Student’s *t*-test and the error bars show SD. Abbreviations: CC, central cell; CCN, central cell nucleus; DSC, degenerated synergid cell; DSN, degenerated synergid nucleus; EC, egg cell; ECN, egg cell nucleus; SC, synergid cell; SCN, synergid cell nucleus. Scale bars, 50 μm.

Given the specific activation of ethylene biosynthesis, signaling, and responses in the persisting synergid cell (Fig. 5a), and the fact that the neighboring zygote and endosperm are seemingly unaffected by gaseous ethylene, we hypothesized that increased ethylene production may trigger cell death exclusively in sensitized synergid cells. To determine the effect of ethylene on female gametophytic cells, we performed ethylene treatment experiments with mature virgin maize ovules. In the mock-treated control, intact synergid nuclei appeared large and round, and the synergid cell status was comparable to untreated materials (n = 119; Fig. 5b). By contrast, in about 40% of ethylene-treated ovules (n = 163), synergid cells exhibited deformed or condensed nuclei (Fig. 5d, e), or showed a highly degenerated status as evidenced by strong autofluorescence (Fig. 5c, e). Striking, the surrounding female gametophytic cells (i.e., egg cell and central cell) were intact and displayed normal nuclei morphologies (Fig. 5c, d). These results imply that synergid cells are especially primed for ethylene-triggered cell death and that the ethylene signaling pathway is highly activated in the persisting synergid cell a few hours before its death.

### PCD of the persistent synergid cell is activated with delay after fertilization

During manual isolation of synergid cells at defined stages during PCD, we observed abrupt removal of the persistent synergid cell after about 25 HAP (Fig. 1l and Supplementary Fig. 1c, d). To investigate the cytological characteristics of this event, we analyzed ovules from maize inbred line B73, as well as two synergid marker lines *pZmES4::ZmES4-GFP* and *pZmSC1::ZmSC1-GFP*, respectively. Before pollination, the synergid cells at the micropylar end of the female gametophyte exhibited clear cell boundaries, with a large and round nucleus situated in the micropylar region of the cell (Fig. 6a). At 24 HAP, the persistent synergid cell still appeared intact and its nucleus displayed largely normal morphology (Fig. 6b). However, after 25 HAP, we observed apparent disappearance of the cell boundary between the persistent synergid cell and the endosperm. Besides, a condensed or disorganized nucleus was located at the micropylar end of the embryo sac, and was eliminated at a slightly later stage (Fig. 6c–e). Detailed analysis of the *pZmSC1::ZmSC1-GFP* marker line further showed that at approximately 25-26 HAP, GFP signals diffused from the persistent synergid cell towards the endosperm, demonstrating the fusion between the persistent synergid cell and the primary endosperm (Fig. 6f, g). This led to a rapid decrease of GFP signals and indicated removal of the persistent synergid cell (Fig. 6f, g and Supplementary Fig. 9c, d). The resulting cytoplasmic continuity between the persistent synergid cell and the endosperm was further supported by observation of manually isolated embryo sacs at 28 HAP showing moderate plasmolysis of a syncytium at the micropylar region of the embryo sac (Supplementary Fig. 9a, b). These observations show that - like in *Arabidopsis*^14^ - the phenomenon of synergid-endosperm fusion (SE fusion) occurs also in the evolutionarily distant cereal crop maize, to quickly remove the persistent synergid cell and thus to terminate pollen tube attraction and other functions of synergid cells.

**Fig. 6.**
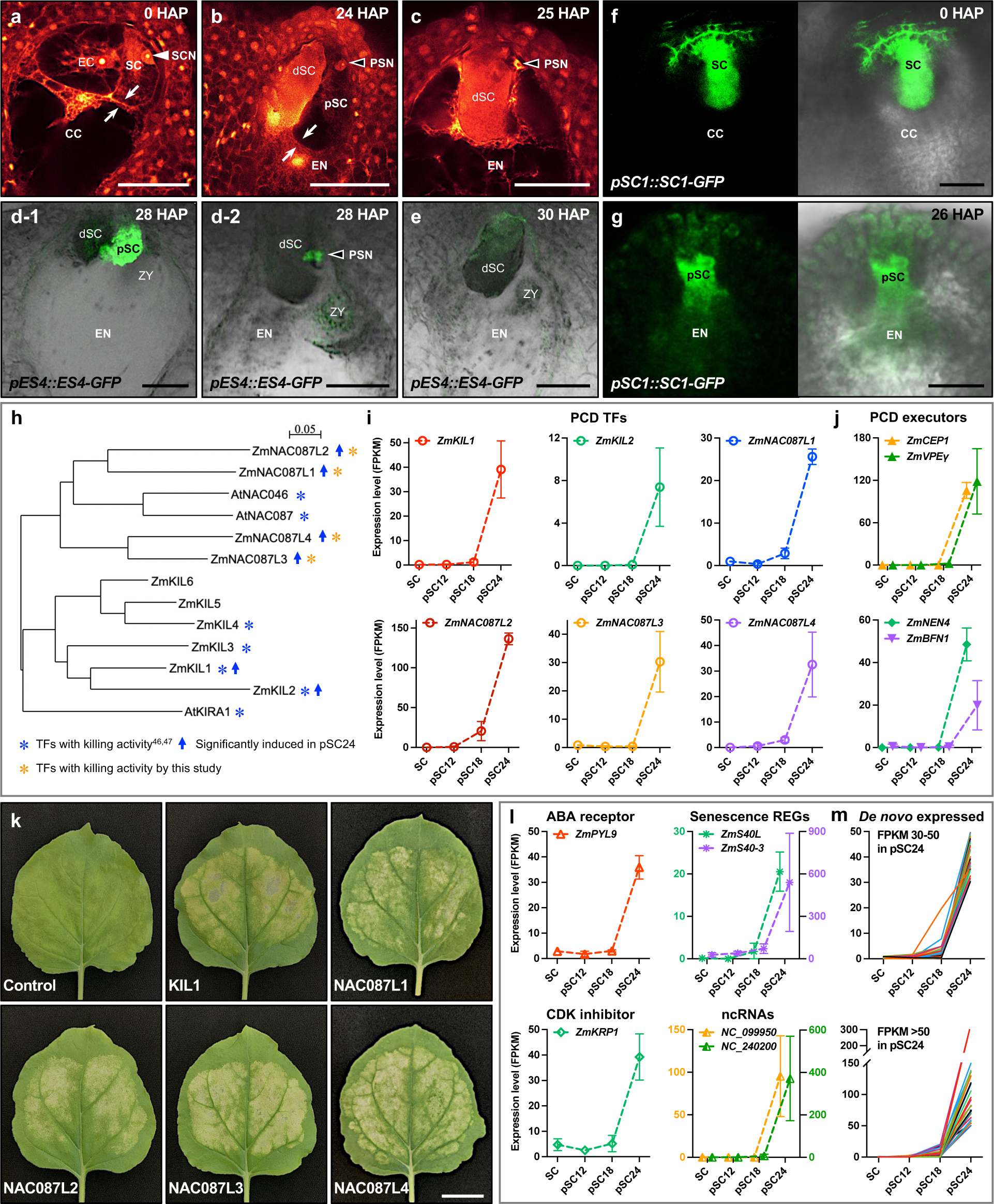
Programmed cell death (PCD) of the persistent synergid cell is activated after successful double fertilization. **a**–**c** CLSM images of embryo sacs before pollination (a) and at 24 HAP (b) and 25 HAP (c), respectively. White arrowhead shows synergid cell nucleus. Black arrowheads point to degenerating persistent synergid nuclei. White arrows indicate cell boundaries. **d**–**g** Fluorescence microscopy to monitor the process of persistent synergid elimination that occurs at about 26-30 HAP. Embryo sacs at indicated time points expressing ES4-GFP and SC1-GFP driven by their endogenous promoters, respectively. The diffusion of GFP signals from persisting synergid cells to the endosperm were observed at 26 HAP (g). **h** Phylogenetic tree of NAC family PCD transcription factors from *Arabidopsis* and their maize homologs expressed during synergid degeneration. **i**,**j** Genes encoding a battery of PCD transcription factors with killing activity and their downstream PCD executors are activated/*de novo* expressed at 18–24 HAP. **k** *Nicotiana benthamiana* leaves infiltrated with NAC family PCD transcription factors that are significantly induced in persisting synergid cells at 24 HAP. **l** Activation patterns of selected maize genes encoding homologs of *Arabidopsis* ABA receptor, CDK inhibitor, and senescence regulators involved in cell death regulation. Two most strongly induced ncRNAs with similar activation patterns are shown. **m** *De novo* expressed unknown genes in the persistent synergid cell showing a PCD-related expression pattern. Abbreviations: CC, central cell; dSC, degenerated synergid cell; EC, egg cell; EN, endosperm; pSC, persistent synergid cell; PSN, persistent synergid nucleus; SC, synergid cell; SCN, synergid cell nucleus; ZY, zygote. Scale bars: 50 μm (a–g) and 2 cm (k), respectively.

Notably, elimination of the persistent synergid has been discussed already before to occur via dPCD^6,7,12,14,17,18^. However, the molecular regulation that ensures the timely activation and execution of PCD processes remained largely elusive. Therefore, we next investigated the expression of maize PCD machinery components homologous to key, well-known *Arabidopsis* PCD transcription factors (TFs) and PCD executors. NAC TFs (<u>N</u>AM: NO APICAL MERISTEM, <u>A</u>TAF1/2: *Arabidopsis thaliana* Activation Factor1 and 2, <u>C</u>UC: CUP-SHAPED COTYLEDON 2) belong to the best-studied TF families in regulating dPCD. ZmKIL1, for example, an ortholog of the *Arabidopsis* NAC TF KIRA1, was recently shown to promote senescence and dPCD in the silk strand base and thus terminate fertility in maize^46^. We found that both, *ZmKIL1* and *ZmKIL2*, were not expressed in synergid cells, but were strongly activated after 18 HAP (Fig. 6h, i and Supplementary Data 8). ANAC087 and ANAC046 have been reported to control dPCD in *Arabidopsis* columella root cap cells^47^. Their homologs in maize (*ZmNAC087L1-4*) exhibited a similar delayed and *de novo* activation pattern in the persisting synergid cell like *ZmKIL1* and *ZmKIL2* (Fig. 6h, i and Supplementary Data 8). We next wondered if the expression of these maize homologs can effectively induce cell death. The leaf transfection assay showed that each of these newly identified candidate dPCD-related NAC TFs (i.e., ZmNAC087L1-4) was sufficient to induce senescence and ectopic cell death (Fig. 6i, k) and therefore represents a novel key molecular player for driving dPCD in plants.

Upon triggering signals and PCD TF activation, PCD execution and corpse clearance are usually initiated^3^. The cysteine protease CEP1, for example, which is transported to the vacuole and transformed into a mature enzyme before rupture of the vacuole, functions as a key executor in tapetal PCD^48^. Vacuolar processing enzymes (VPEs), major vacuolar proteases with caspase-like activity, can activate other vacuolar hydrolases, and thus are in control of tonoplast rupture and vacuole collapse^2,49^. VPEs translocate to the vacuole through the autophagy pathway^50^, and autophagy is functionally implicated in corpse clearance during dPCD^51,52^. We searched from maize homologs and found a dramatic *de novo* activation of *ZmCEP1* and *ZmVPEγ* in persistent synergid cells at 24 HAP (Fig. 6j), being consistent with a clear activation of autophagic processes for PCD execution (Fig. 5g, h). Besides vacuolar proteases, three putative NAC-dependent nucleases (ZmENDO1, ZmNEN1/4) were simultaneously *de novo* induced during PCD execution and corpse clearance (Fig. 6j and Supplementary Data 8). Taken together, these results show that autophagy and key vacuolar cysteine proteases have been activated for vacuole-related corpse clearance before persistent synergid elimination. Moreover, morphological disorganization of the persistent synergid cell nucleus - another characteristic of PCD - was observed demonstrating that PCD precedes SE-fusion and thus ultimately synergid cell elimination.

Besides the canonical PCD TFs and PCD executors, a subset of genes involved in PCD regulation were also strongly activated. These include orthologs of ABA receptor *PYL9* and CDK inhibitor *KRP1*, senescence regulators, as well as highly induced ncRNAs (Fig. 6l). In total, among the 2,727 genes showing significant upregulation after fertilization at 24 HAP, 379 genes were *de novo* expressed (only genes with an FPKM > 5 were considered; Supplementary Data 9), including above described PCD regulators, TFs and executors. Notably, nearly all of them exhibited a PCD-related expression pattern, that is, not expressed before 18 HAP (∼10 HAF) or slightly induced at 18 HAP, but dramatically activated shortly before persistent synergid elimination (Fig. 6m). This includes also many so far uncharacterized genes, representing very likely components of the molecular machinery responsible for driving dPCD in plants.

## Conclusions

In conclusion, our data reveal a high-resolution atlas of timely activation and execution of dPCD, and additionally provides a rich resource for future studies investigating novel dPCD regulators. Studying the elimination of the persistent synergid cell is uniques as it allows to access cells undergoing dPCD at highly precise timing as induction of dPCD is triggered in this case by fertilization. Cells disintegrate during about 18 h after fertilization due to the activation of genes encoding executors and degraders shortly before cell elimination. In addition to the GRN controlled by MYB98 required for synergid functions before fertilization (Fig. 7a), we show that apparently further networks controlled by HB transcription factors are downregulated after dPCD induction. The overall decrease of most highly expressed synergid-specific transcription factors and their targets appears to go hand-in-hand with the loss of synergid cell identity and functions. Exploring activated signaling pathways during synergid PCD, we demonstrate how fertilization-induced synergid-derived autocrine peptide signal RALF12 specifically triggers mitochondrial ROS production and accumulation at the filiform apparatus region, accompanied by downregulation of mitochondrial antioxidant enzymes. This seems to mediate oxidative stress and activates dPCD in persisting synergid cells. In addition, both, ethylene production and key ethylene signaling components are specifically activated shortly after successful fertilization, which further promotes the process of sensitized synergid cell death.

**Fig. 7.**
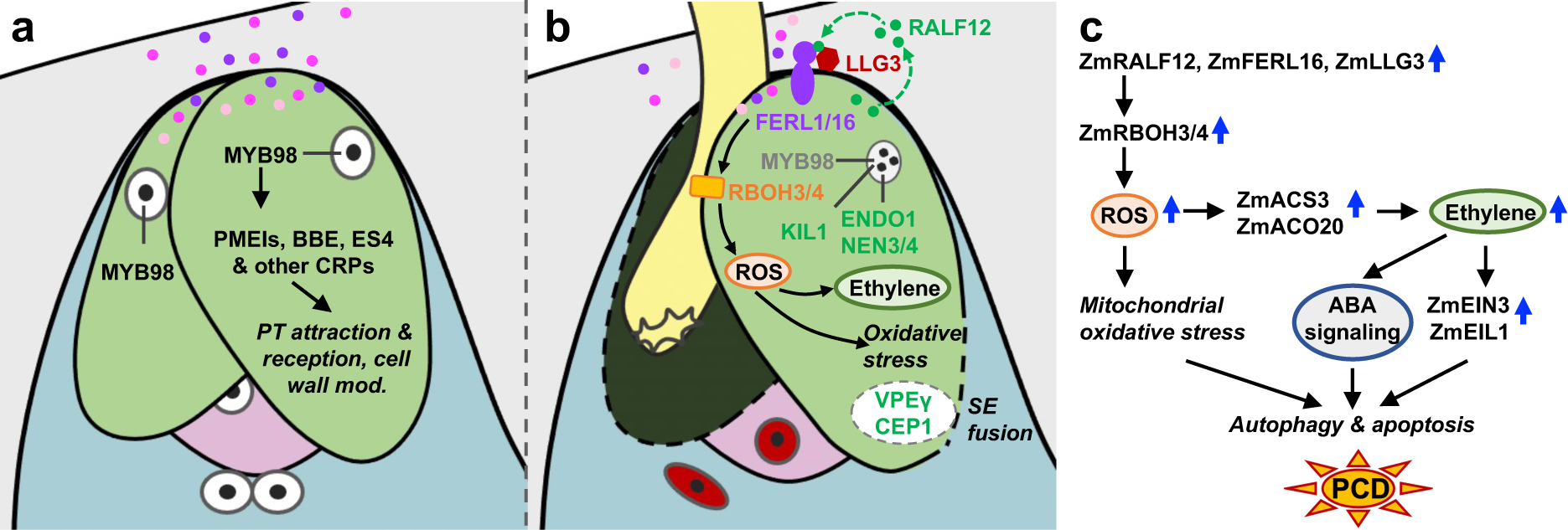
Model of ZmMYB98 controlled synergid cell functions and subsequent ZmRALF12 induced persistent synergid PCD after fertilization. **a** Before fertilization, ZmMYB98 positively regulates most highly expressed synergid-specific genes required for pollen tube attraction and reception as well as cell wall modification. **b** After successful double fertilization, ZmMYB98 and its associated gene regulatory network (GRN) are downregulated, while a GRN is activated leading to RALF12 signaling culminating in high levels of mitochondrial ROS accumulation at the filiform apparatus region, oxidative stress, mitophagy, and synergid apoptosis. Activated ethylene biosynthesis further promotes sensitized synergid cell death. RALF12-mediated mitochondrial oxidative stress and ethylene biosynthesis can individually trigger synergid cell death, while they work synergistically to highly efficiently activate PCD TFs and PCD executors for persistent synergid nucleus and whole-cell elimination, in a few hours after successful double fertilization. Vacuole rupture as well as cell intercalation mediated by VPEγ and CEP1 likely contribute to synergid-endosperm (SE) fusion. **c** Schematic model of the GRN leading to persistent synergid PCD.

Moreover, this work clarifies that persistent synergid cells indeed undergo PCD before SE fusion and that this fusion event, which has so far only been described in *Arabidopsis*^14^, is likely conserved in flowering plants. Notably, we show that being consistent with the specific and significant activation of *ACOs* in persisting synergid cells, the gaseous plant hormone ethylene is capable to induce death specifically of synergid cells, but not of the surrounding female gametes. This is reflected by the presence of the sensitive ethylene perception, signaling, and response machinery in synergid cells, but not in the surrounding cells.

As further summarized in our working model (Fig. 7b, c), this study indicates how mitochondrial ROS-induced dPCD, marked by the onset of oxidative stress, works synergistically with activated ethylene production and signaling to ensure timely dPCD execution and corpse clearance shortly before SE fusion. It will now be exciting to find out whether misexpression of RALF12 is sufficient to trigger PCD also in other cell types and whether RALF12-triggered ROS production in synergid cells is consertvedf among plants. Functional studies are now necessary to elucidate the roles of previously undescribed dPCD-activation-related genes to better understand the molecular machinery responsible for driving PCD during development.

## Methods

### Plant material, transgenic plants and growth conditions

Maize (*Zea mays*) inbred line B73 was used for manual cell isolation and cell-specific RNA-seq. Transgenic line *pZmES4::ZmES4-GFP* was described previously^20^ and *pZmSC1::ZmSC1-GFP* as well as *pZmSC3::ZmSC3-GFP* transgenic lines were generated in the B104 background. To generate GFP reporter lines, gene promoter sequences (around 3.2 kbp of *ZmSC1* and 2.6 kbp of *ZmSC3*) and their respective coding sequences were amplified and cloned into the vector pTF101.1^53^ upstream of GFP. Constructs were transformed into B104 via *Agrobacterium*-mediated transformation by Edgene Biotech company. Primer sequences used in this study are shown in Supplementary Data 10.

Plants were cultivated in the greenhouses at 26°C under illumination of 24,000 lux with 16 h light/8 h dark cycles and a relative humidity of 60%.

### Isolation of synergid cells with precise timing

Synergid cell stages are defined according to the precise time after pollination/fertilization. To isolate synergid cells at defined stages, ovules in the middle part of maize cobs were collected and sliced manually. Ovular sections containing embryo sacs were treated for 10 min with 1% cellulase R10 and 0.5% macerozyme R10 (Yakult Pharmaceutical Ind. Co. Ltd.) dissolved in 11% mannitol and 0.058% MES pH 5.8. Next, samples were extensively washed in the same solution without cell wall degrading enzyme. Microdissection of synergid/persistent synergid cells was performed as previously described^26,54^. All cells isolated from ovules were washed three times in the same solution, individually transferred to 0.5 mL RNA/DNA LoBind microcentrifuge tubes (Eppendorf), immediately frozen in liquid nitrogen, and stored at −80°C for mRNA extraction.

### cDNA preparation and library construction for RNA-seq

RNA extraction and cDNA preparation were performed according to our previous protocol with minor modifications^21^. Briefly, mRNA was isolated from cell samples using the Dynabeads mRNA DIRECT^TM^ Micro Kit (Life Technologies). cDNA synthesis and amplification were performed using a SMART-Seq v4 Ultra Low Input RNA Kit for sequencing (Clontech). cDNA was purified using an Agencourt AMPure purification kit (Beckman Coulter). After purification and quantification, cDNA was used for library construction. RNA-seq libraries were prepared using a NEBNext Ultra DNA Library Prep Kit for Illumina (New England Biolabs) according to the manufacturer’s instructions, and sequenced on an Illumina Novaseq platform using paired-end 150 bp flow cells.

### RNA-seq data analysis

Clean reads of samples were prepared using Cutadapt version 1.15^55^ and an inhouse perl script. Clean reads were mapped to the maize genome (AGPv5, Zm-B73-REFERENCE-NAM-5.0, http://ftp.ensemblgenomes.org/pub/plants/release-51/fasta/zea_mays/dna/) using Hisat2 version 2.0.5^56^. Gene expression levels were quantified as FPKM using featureCounts version 1.5.0-p3^57^. Differential expression analysis of stage-to-stage comparisons (three biological replicates per stage) was performed using DESeq2^58^. Genes with a |log_2_FC| > 1 and false discovery rate (FDR) < 0.05 were considered significant differentially expressed.

### Dual-luciferase reporter assay

A dual-luciferase reporter assay was performed to analyze promoter activities. To generate reporter constructs, promoters (around 2 kbp) of the 19 most highly expressed synergid-specific genes were amplified and cloned upstream of the *LUC* gene in the reporter vector pGreen II 0800-LUC^59^. The full-length coding sequence of *ZmMYB98* was cloned into pGreen II 62-SK^59^ plasmid downstream of the cauliflower mosaic virus (CaMV) 35S promoter to generate an effector vector. *Agrobacterium* cells containing the reporter and effector constructs were mixed and infiltrated into tobacco leaves. After incubating for 72 h, infected areas of the leaves were collected to measure the activity of firefly luciferase (LUC) and renilla luciferase (REN) using a Dual Luciferase Reporter Gene Assay Kit (Beyotime Biotech). Three biological replicates were performed for each experiment. Primer sequences used are listed in Supplementary Data 10.

### ROS staining and quantification

For ROS stainings, micropylar ovule regions containing embryo sacs or manually isolated synergid cells were dissected before pollination and at 20 HAP, respectively. A Reactive Oxygen Species Assay Kit (Beyotime Biotech) was used to detect intracellular reactive oxygen species. Samples were immersed in ROS staining solution (7.5 μM DCFH-DA in 11% mannitol solution, pH 5.8) in the dark for 10 min, and then washed three times with 11% mannitol pH 5.8. Imaging was carried out using a Leica confocal laser scanning microscope (CLSM) SP8 with an excitation wavelength of 488 nm. Laser intensities for all comparative samples were exactly the same within one experiment. ImageJ^60^ was used to determine fluorescence intensity.

For ROS and Mito-Tracker double staining, cells were incubated in double staining solution (7.5 μM DCFH-DA and 20 nM Mito-Tracker Red CMXRos [Beyotime Biotech] in 11% mannitol solution, pH 5.8) in the dark for 10 min, extensively washed in 11% mannitol solution, and then imaged using a CLSM SP8 with an excitation wavelength of 488 nm (ROS) and 552 nm (Mito-Tracker). The relative fluorescent intensity of cells was measured using the LAS-X software (Leica).

### RALF peptide preparation and treatments

*ZmRALF1*, *ZmRALF2*, and *ZmRALF12* coding sequences^24^ without signal peptides were amplified and cloned into the pET-32a (Novagen) vector, respectively. The generated vector was transformed into *E. coli* strain BL21 (DE3). Recombinant ZmRALF1, ZmRALF2, and ZmRALF12 (RALF) were expressed according to the PET system instructions (Novagen), and purified with TALON Metal Affinity Resin (Clontech) according to the manufacturer’s instructions.

Synergid cells from mature virgin ovules were manually isolated and collected in 11% mannitol solution. For RALF treatment, cells were incubated in solution of each 4 μM RALF peptide in 11% mannitol solution for 30 min. Cell samples were extensively washed after incubation. ROS staining procedures were immediately performed after treatment as described above.

### MDC staining

Synergid cells and persistent synergid cells at 24 HAP were manually isolated and collected in 11% mannitol solution. Autophagic structures in cells were visualized by fluorescent monodansylcadaverine (MDC) staining using an Autophagy Staining Assay Kit with MDC (Beyotime Biotech) following the manufacturer’s instructions. Images were captured with a Leica TCS SP8 CLSM with 405 nm excitation.

### Ethylene treatment and analysis

For ethylene treatment, the middle parts of unpollinated cobs were placed in 100 ml sealed glass bottles to incubate with gaseous ethylene released from ethephon (Aladin Biotech). 0.1 g ethephon was dissolved in 1 ml sterile pure water in a small glass dish beside dissected cobs. After 30 h, cobs were collected and treated for CLSM analysis according to a protocol described previously with some modifications^61^. Ovules were dissected with two longitudinal sections along the silk axis. Ovule pieces containing embryo sacs were fixed in a solution of 4% glutaraldehyde in 12.5 mM cacodylate (pH 6.9) for 2 h at room temperature and then at 4°C overnight. Following fixation, samples were dehydrated by sequential treatment with 10%, 20%, 40%, 60%, 85%, 95%, and 100% ethanol for 30 min at each step. After dehydration, samples were cleared in a 2:1 mixture of benzyl benzoate: benzyl alcohol for 20 min. Cleared sections were analyzed using a Leica TCS SP8 CLSM with an excitation wavelength of 488 nm.

### Confocal microscopy analysis of synergid degeneration processes

To examine synergid cell morphology and its removal at defined developmental stages, ovules were collected and manually dissected along the silk axis. Ovular sections containing embryo sacs were fixed and analyzed as described above for ethylene-treated samples. To visualize GFP and SE fusion dynamics, *pZmES4::ZmES4-GFP* and *pZmSC1::ZmSC1-GFP* cobs were harvested at indicated stages, ovules were dissected and kept in 11% mannitol solution. Microscopy was performed on the Leica TCS SP8 CLSM with 488nm excitation. Laser intensities for all comparative samples were exactly the same within one experiment.

## Acknowledgments

We would like to thank Mengxiang Sun and Xiongbo Peng (Wuhan University) for assistance in teaching manual cell isolation techniques. Wenliang Xu (Central China Normal University) is acknowledged for help with the dual-luciferase assay system and Armin Hildebrand (University of Regensburg) for plant care. This work was supported by grants from the National Natural Science

Foundation of China (31970342 to J.C.), the Foundation of Hubei Hongshan Laboratory (2022hszd017 to J.C.), the Ten Thousand Talents Program for Young Talents (to J.C.), and the Alexander von Humboldt Foundation (to X.Z. and T.D.).

## Author contributions

J.C. and T.D. conceptualized the study, designed the research plan, interpreted the results and wrote the manuscript with input from all authors. H.W., J.W. and J.C. collected cell samples, J.C. analyzed transcriptomic data with support from T.D., L.H. and S. H.; J.W. performed ROS staining experiments, H.W. and J.W. dual-luciferase assays, RALF and ethylene treatments, while X.Z., W.Q, H.F. S.W. and H.W. conducted CLSM observation of transgenic plants. All authors contributed to data collection and discussion, presentation and finalizing the manuscript.

## Additional information

### Supplementary Information

Supplementary Fig. 1: Timing of synergid cell death during the fertilization process in maize.

Supplementary Fig. 2: RNA-seq analysis at precise synergid degenerating stages.

Supplementary Fig. 3: The major transcriptional activation and repression wave in the persistent synergid cell occurs about 24 HAP.

Supplementary Fig. 4: *In planta* activation of synergid-specific gene promoters by ZmMYB98 and secretion of gene products.

Supplementary Fig. 5: RALF12 triggers high levels of granular ROS accumulation at the filiform apparatus region in a time-and concentration-dependent manner.

Supplementary Fig. 6: Peroxisomal function-related genes are significantly activated indicating an oxidative stress status in synergids after 18 HAP.

Supplementary Fig. 7: Genes encoding Ca^2+^ channels, Ca^2+^ sensors, and Ca^2+^ signaling components involved in senescence and cell death regulation are activated in synergids after 18 HAP.

Supplementary Fig. 8: Autophagy-related genes are activated with delay after fertilization in synergids for corpse clearance.

Supplementary Fig. 9: Elimination of the persistent synergid cell through synergid-endosperm (SE) fusion at the end of PCD.

### Supplementary Data

Supplementary Data 1: Summary of cell-and stage-specific RNA-Seq library statistics and read mapping results.

Supplementary Data 2: Expression level of each gene in each cell degenerating stage.

Supplementary Data 3: Subcellular distribution of most abundant gene products in different cells.

Supplementary Data 4: The most highly expressed genes in synergid cells encode especially synergid-specific secreted peptides and secreted cell wall modifiers.

Supplementary Data 5: The most strongly expressed TFs are synergid-specific and are substantially decreased during synergid degeneration.

Supplementary Data 6: Downregulation of most highly expressed genes during synergid degeneration.

Supplementary Data 7: The major transcriptional activation and repression wave occurs about 24 HAP (timing of successful fertilization).

Supplementary Data 8: Activation of ROS production, oxidative stress response, ethylene biosynthesis and signaling, and PCD regulation during synergid PCD.

Supplementary Data 9: De nove expressed genes all exhibite PCD-related expression patterns.

Supplementary Data 10: Primers used in the present study.

## References

1 Daneva, A., Gao, Z., Van Durme, M. & Nowack, M. K. Functions and regulation of programmed cell death in plant development. Annu. Rev. Cell. Dev. Biol. 32, 441–468 (2016).

2 Dickman, M., Williams, B., Li, Y., de Figueiredo, P. & Wolpert, T. Reassessing apoptosis in plants. Nat. Plants 3, 773–779 (2017).

3 Huysmans, M., Lema, A. S., Coll, N. S. & Nowack, M. K. Dying two deaths - programmed cell death regulation in development and disease. Curr. Opin. Plant Biol. 35, 37–44 (2017).

4 Xie, F., Vahldick, H., Lin, Z. & Nowack, M. K. Killing me softly - Programmed cell death in plant reproduction from sporogenesis to fertilization. Curr. Opin. Plant Biol. 69, 102271 (2022).

5 Leydon, A. R. et al. Pollen tube discharge completes the process of synergid degeneration that is initiated by pollen tube-synergid interaction in *Arabidopsis*. Plant Physiol. 169, 485–496 (2015).

6 Völz, R., Harris, W., Hirt, H. & Lee, Y. H. ROS homeostasis mediated by MPK4 and SUMM2 determines synergid cell death. Nat. Commun. 13, 1746 (2022).

7 Hater, F., Nakel, T. & Groß-Hardt, R. Reproductive multitasking: the female gametophyte. Annu. Rev. Plant Biol. 71, 517–546 (2020).

8 Denninger, P. et al. Male-female communication triggers calcium signatures during fertilization in *Arabidopsis*. Nat. Commun. 5, 4645 (2014).

9 Ngo, Q. A., Vogler, H., Lituiev, D. S., Nestorova, A. & Grossniklaus, U. A calcium dialog mediated by the FERONIA signal transduction pathway controls plant sperm delivery. Dev. Cell 29, 491–500 (2014).

10 Duan, Q. et al. Reactive oxygen species mediate pollen tube rupture to release sperm for fertilization in *Arabidopsis*. Nat. Commun. 5, 3129 (2014).

11 Kasahara, R. D. et al. Fertilization recovery after defective sperm cell release in *Arabidopsis*. Curr. Biol. 22, 1084–1089 (2012).

12 Johnson, M. A., Harper, J. F. & Palanivelu, R. A fruitful journey: pollen tube navigation from germination to fertilization. Annu. Rev. Plant Biol. 70, 809–837 (2019).

13 Wong, J. L. & Wessel, G. M. Defending the zygote: search for the ancestral animal block to polyspermy. Curr. Top. Dev. Biol. 72, 1–151 (2006).

14 Maruyama, D. et al. Rapid elimination of the persistent synergid through a cell fusion mechanism. Cell 161, 907–918 (2015).

15 Maruyama, D. & Higashiyama, T. The end of temptation: the elimination of persistent synergid cell identity. Curr. Opin. Plant Biol. 34, 122–126 (2016).

16 Sandaklie-Nikolova, L., Palanivelu, R., King, E. J., Copenhaver, G. P. & Drews, G. N. Synergid cell death in Arabidopsis is triggered following direct interaction with the pollen tube. Plant Physiol. 144, 1753–1762 (2007).

17 Völz, R., Heydlauff, J., Ripper, D., von Lyncker, L. & Groß-Hardt, R. Ethylene signaling is required for synergid degeneration and the establishment of a pollen tube block. Dev. Cell 25, 310–316 (2013).

18 Li, W. et al. Lack of ethylene does not affect reproductive success and synergid cell death in *Arabidopsis*. Mol. Plant 15, 354–362 (2022).

19 Mendes, M. A. et al. Live and let die: a REM complex promotes fertilization through synergid cell death in *Arabidopsis*. Development 143, 2780–2790 (2016).

20 Amien, S. et al. Defensin-like ZmES4 mediates pollen tube burst in maize via opening of the potassium channel KZM1. PLoS Biol. 8, e1000388 (2010).

21 Chen, J. et al. Zygotic genome activation occurs shortly after fertilization in maize. Plant Cell 29, 2106–2125 (2017).

22 Escobar-Restrepo, J. M. et al. The FERONIA receptor-like kinase mediates male-female interactions during pollen tube reception. Science 317, 656–660 (2007).

23 Liu, X. et al. The role of LORELEI in pollen tube reception at the interface of the synergid cell and pollen tube requires the modified eight-cysteine motif and the receptor-like kinase FERONIA. Plant Cell 28, 1035–1052 (2016).

24 Zhou, L. Z. et al. The RALF signaling pathway regulates cell wall integrity during pollen tube growth in maize. Plant Cell, DOI: 10.1093/plcell/koad1324 (2023).

25 Wu, J. J. et al. Mitochondrial GCD1 dysfunction reveals reciprocal cell-to-cell signaling during the maturation of *Arabidopsis* female gametes. Dev. Cell 23, 1043–1058 (2012).

26 Cordts, S., et al. *ZmES* genes encode peptides with structural homology to defensins and are specifically expressed in the female gametophyte of maize. Plant J. 25, 103–114 (2001).

27 Zhong, S. & Qu, L. J. Peptide/receptor-like kinase-mediated signaling involved in male-female interactions. Curr. Opin. Plant Biol. 51, 7–14 (2019).

28 Higashiyama, T. & Takeuchi, H. The mechanism and key molecules involved in pollen tube guidance. Annu. Rev. Plant Biol. 66, 393–413 (2015).

29 Dresselhaus, T., Sprunck, S. & Wessel, G. M. Fertilization mechanisms in flowering plants. Curr. Biol. 26, R125–139 (2016).

30 Punwani, J. A., Rabiger, D. S. & Drews, G. N. MYB98 positively regulates a battery of synergid-expressed genes encoding filiform apparatus localized proteins. Plant Cell 19, 2557–2568 (2007).

31 Zhou, L. Z. & Dresselhaus, T. Multiple roles of ROS in flowering plant reproduction. Adv. Bot. Res. 105, 139–176 (2022).

32 Mittler, R. ROS are good. Trends Plant Sci. 22, 11–19 (2017).

33 Mittler, R., Zandalinas, S. I., Fichman, Y. & Van Breusegem, F. Reactive oxygen species signalling in plant stress responses. Nat. Rev. Mol. Cell Biol. 23, 663–679 (2022).

34 Ge, Z., Cheung, A. Y. & Qu, L. J. Pollen tube integrity regulation in flowering plants: insights from molecular assemblies on the pollen tube surface. New Phytol. 222, 687–693 (2019).

35 Liu, C. et al. Pollen PCP-B peptides unlock a stigma peptide-receptor kinase gating mechanism for pollination. Science 372, 171–175 (2021).

36 Ermak, G. & Davies, K. J. Calcium and oxidative stress: from cell signaling to cell death. Mol. Immunol. 38, 713–721 (2002).

37 Orrenius, S., Gogvadze, V. & Zhivotovsky, B. Mitochondrial oxidative stress: implications for cell death. Annu. Rev. Pharmacol. Toxicol. 47, 143–183 (2007).

38 Andrabi, S. S., Yang, J., Gao, Y., Kuang, Y. & Labhasetwar, V. Nanoparticles with antioxidant enzymes protect injured spinal cord from neuronal cell apoptosis by attenuating mitochondrial dysfunction. J Control Release 317, 300–311 (2020).

39 Broda, M., Millar, A. H. & Van Aken, O. Mitophagy: a mechanism for plant growth and survival. Trends Plant Sci. 23, 434–450 (2018).

40 Xu, E., Vaahtera, L. & Brosché, M. Roles of defense hormones in the regulation of ozone-induced changes in gene expression and cell death. Mol. Plant 8, 1776–1794 (2015).

41 Waszczak, C., Carmody, M. & Kangasjärvi, J. Reactive oxygen species in plant signaling. Annu. Rev. Plant Biol. 69, 209–236 (2018).

42 Heydlauff, J. et al. Dual and opposing roles of EIN3 reveal a generation conflict during seed growth. Mol. Plant 15, 363–371 (2022).

43 Adams, D. O. & Yang, S. F. Ethylene biosynthesis: Identification of 1-aminocyclopropane-1-carboxylic acid as an intermediate in the conversion of methionine to ethylene. Proc. Natl. Acad. Sci. USA 76, 170–174 (1979).

44 Pattyn, J., Vaughan-Hirsch, J. & Van de Poel, B. The regulation of ethylene biosynthesis: a complex multilevel control circuitry. New Phytol. 229, 770–782 (2021).

45 Chao, Q. et al. Activation of the ethylene gas response pathway in *Arabidopsis* by the nuclear protein ETHYLENE-INSENSITIVE3 and related proteins. Cell 89, 1133–1144 (1997).

46 Šimášková, M. R. et al. KIL1 terminates fertility in maize by controlling silk senescence. Plant Cell 34, 2852–2870 (2022).

47 Huysmans, M. et al. ANAC087 and ANAC046 control distinct aspects of programmed cell death in the *Arabidopsis* columella and lateral root cap. Plant Cell 30, 2197–2213 (2018).

48 Zhang, D. et al. The cysteine protease CEP1, a key executor involved in tapetal programmed cell death, regulates pollen development in *Arabidopsis*. Plant Cell 26, 2939–2961 (2014).

49 Wleklik, K. & Borek, S. Vacuolar processing enzymes in plant programmed cell death and autophagy. Int. J. Mol. Sci. 24 (2023).

50 Teper-Bamnolker, P. et al. Vacuolar processing enzyme translocates to the vacuole through the autophagy pathway to induce programmed cell death. Autophagy 17, 3109–3123 (2021).

51 Minina, E. A., Bozhkov, P. V. & Hofius, D. Autophagy as initiator or executioner of cell death. Trends Plant Sci. 19, 692–697 (2014).

52 Feng, Q., De Rycke, R., Dagdas, Y. & Nowack, M. K. Autophagy promotes programmed cell death and corpse clearance in specific cell types of the *Arabidopsis* root cap. Curr. Biol. 32, 2110–2119.e2113 (2022).

53 Paz, M. M. et al. Assessment of conditions affecting *Agrobacterium*-mediated soybean transformation using the cotyledonary node explant. Euphytica 136, 167–179 (2004).

54 Kranz, E., Bautor, J. & Lörz, H. In vitro fertilization of single, isolated gametes of maize mediated by electrofusion. Sex. Plant Reprod. 4, 12–16 (1991).

55 Martin, M. Cutadapt removes adapter sequences from high-throughput sequencing reads. EMBnet J. 17, 10–12 (2011).

56 Mortazavi, A., Williams, B. A., McCue, K., Schaeffer, L. & Wold, B. Mapping and quantifying mammalian transcriptomes by RNA-Seq. Nat. Methods 5, 621–628 (2008).

57 Liao, Y., Smyth, G. K. & Shi, W. featureCounts: an efficient general purpose program for assigning sequence reads to genomic features. Bioinformatics 30, 923–930 (2014).

58 Love, M. I., Huber, W. & Anders, S. Moderated estimation of fold change and dispersion for RNA-seq data with DESeq2. Genome Biol. 15, 1–21 (2014).

59 Hellens, R. P. et al. Transient expression vectors for functional genomics, quantification of promoter activity and RNA silencing in plants. Plant methods 1, 1–14 (2005).

60 Collins, T. J. ImageJ for microscopy. BioTechniques 43, S25–S30 (2007).

61 Christensen, C. A., King, E. J., Jordan, J. R. & Drews, G. Megagametogenesis in *Arabidopsis* wild type and the *Gf* mutant. Sex. Plant Reprod. 10, 49–64 (1997).

