## Supplemental Figures S1-S9 for "Fertilization-induced synergid cell death by RALF12-triggered ROS production and ethylene signaling"

### Supplementary Information

| | |
|---|---|
| Supplementary Fig. 1: | Timing of synergid cell death during the fertilization process in maize. |
| Supplementary Fig. 2: | RNA-seq analysis at precise synergid degenerating stages. |
| Supplementary Fig. 3: | The major transcriptional activation and repression wave in the persistent synergid cell occurs about 24 HAP. |
| Supplementary Fig. 4: | *In planta* activation of synergid-specific gene promoters by ZmMYB98 and secretion of gene products. |
| Supplementary Fig. 5: | RALF12 triggers high levels of granular ROS accumulation at the filiform apparatus region in a time- and concentration-dependent manner. |
| Supplementary Fig. 6: | Peroxisomal function-related genes are significantly activated indicating an oxidative stress status in synergids after 18 HAP. |
| Supplementary Fig. 7: | Genes encoding $Ca^{2+}$ channels, $Ca^{2+}$ sensors, and $Ca^{2+}$ signaling components involved in senescence and cell death regulation are activated in synergids after 18 HAP. |
| Supplementary Fig. 8: | Autophagy-related genes are activated with delay after fertilization in synergids for corpse clearance. |
| Supplementary Fig. 9: | Elimination of the persistent synergid cell through synergid-endosperm (SE) fusion at the end of PCD. |

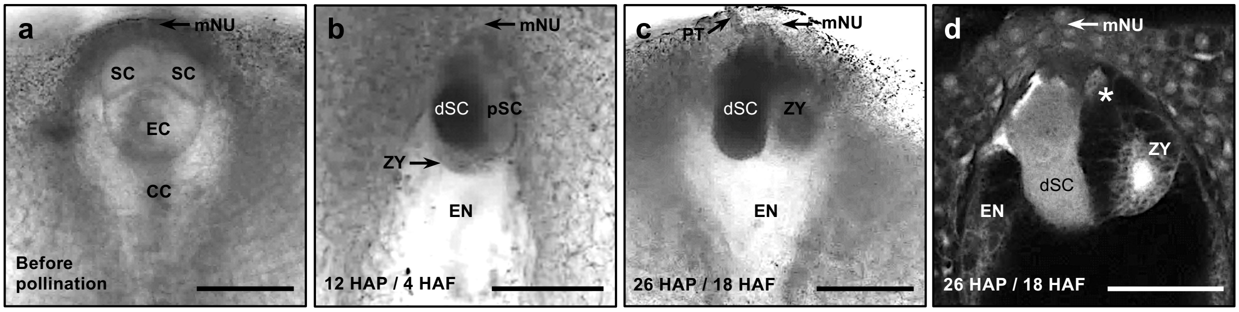


**Supplementary Fig. 1 Timing of synergid cell death during the fertilization process in maize. a**,**b** Bright-field micrographs showing the embedded female gametophyte (embryo sac) containing two synergid cells before fertilization and at 12 HAP (this corresponds to about 4 HAF), respectively. The receptive synergid invaded by a pollen tube appears dark beside the persistent synergid cell. **c**,**d** Representative bright-field (c) and CLSM (d) images of embryo sacs at 26 HAP (about 18 HAF). The persistent synergid is no longer visible, leaving the degenerated receptive synergid and the zygote at the micropylar region of the embryo sac. Asterisk marks residues of the disappeared persistent synergid cell. Abbreviations: CC, central cell; dSC, degenerated synergid cell; EC, egg cell; EN, endosperm; HAP, hours after pollination; HAF, hours after fertilization; mNU, micropylar nucellus; pSC, persistent synergid cell; PT, pollen tube; SC, Synergid cell; ZY, zygote. Scale bars, 50 μm.

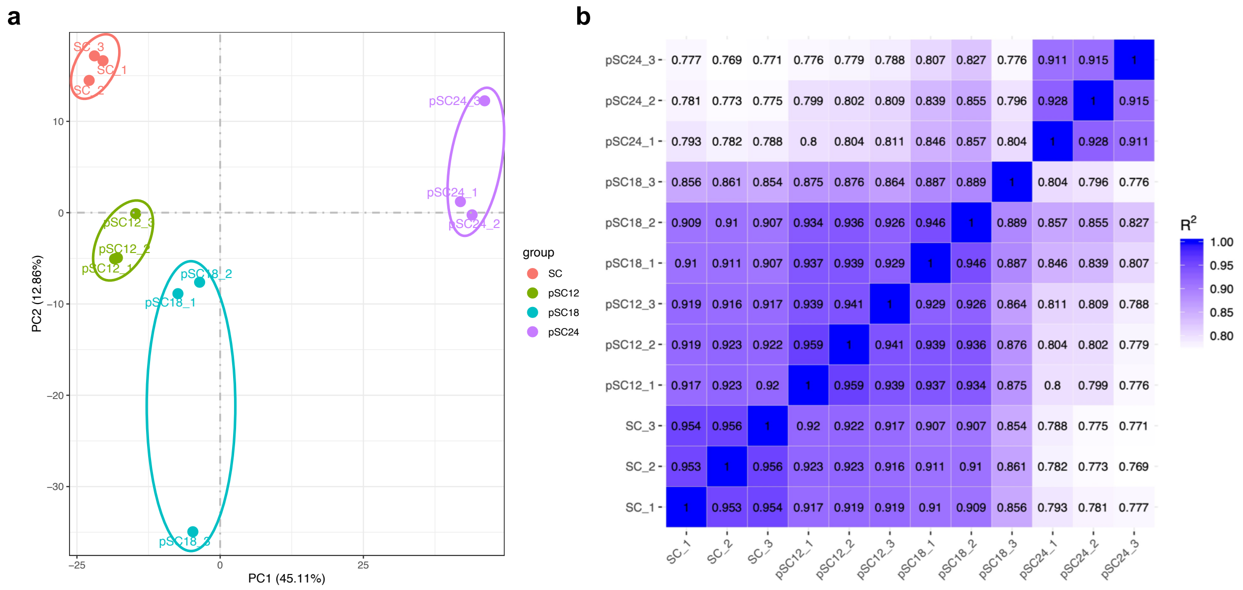


**Supplementary Fig. 2 RNA-seq analysis at precise synergid degenerating stages. a** Principal component analysis (PCA) of synergid cell RNA-seq data at indicated stages. **b** RNA-seq data of the same synergid stage are highly correlated. The correlation coefficients between different samples are calculated according to Pearson's correlation coefficient method. Abbreviations: SC, synergid cell before pollination; pSC12, persistent synergid cell at 12 hours after pollination (HAP); pSC18, persistent synergid cell at 18 HAP; pSC24, persistent synergid cell at 24 HAP.

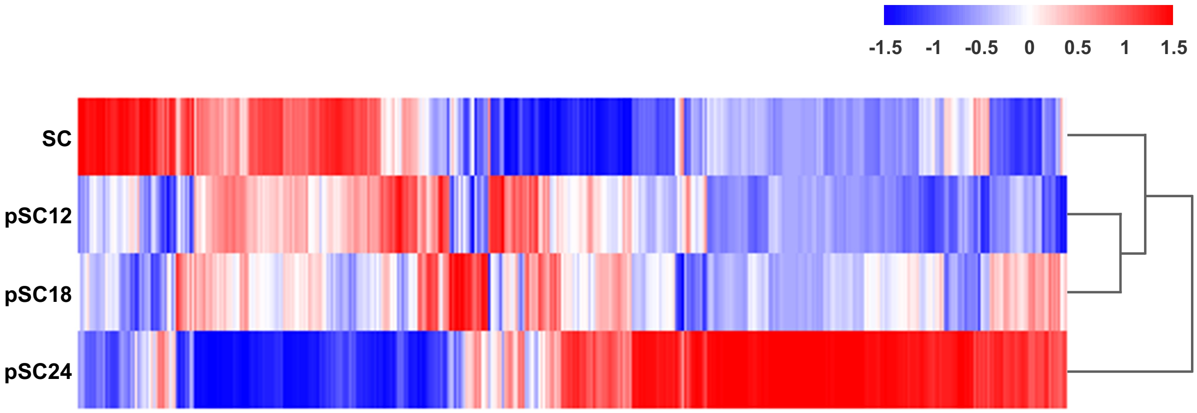


**Supplementary Fig. 3 The major transcriptional activation and repression wave in the persistent synergid cell occurs about 24 HAP.** Heatmap showing the expression pattern of differentially expressed genes (DEGs) in synergid cells across precise stages during fertilization and synergid degeneration. Genes with abs($log_2$FC) > 1 and adjusted $P < 0.05$ in at least one synergid cell degenerating stage comparison are included.

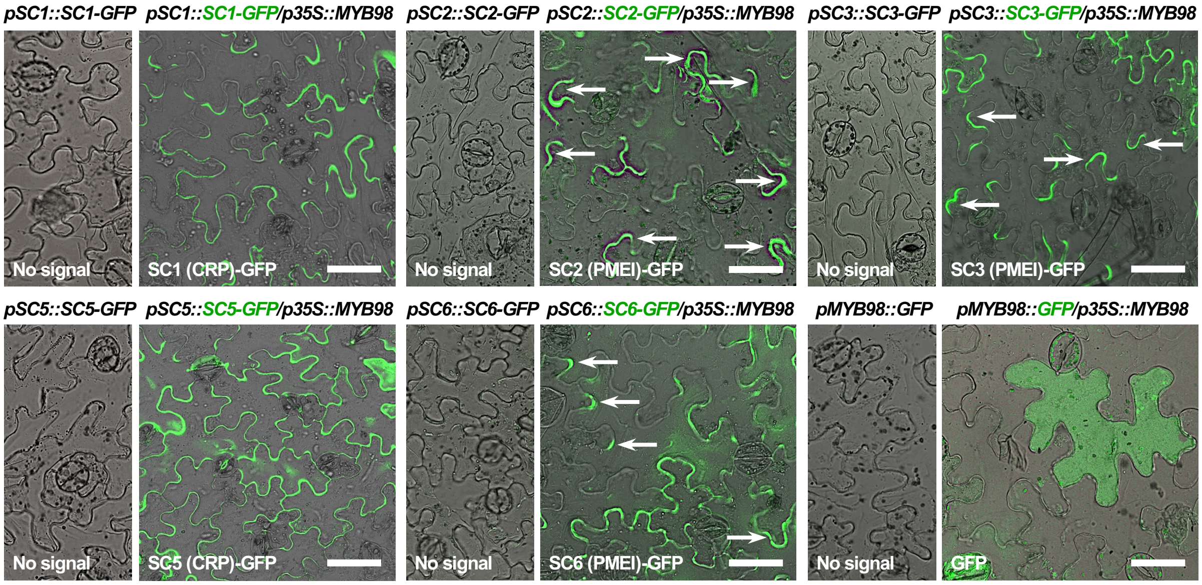


**Supplementary Fig. 4 *In planta* activation of synergid-specific gene promoters by ZmMYB98 and secretion of gene products.** *N. benthamiana* leaves were infiltrated with indicated constructs. *pMYB98::GFP* was used as a control. The most highly expressed synergid-specific gene products were secreted to the extracellular region. Arrows indicate the strongest signals at the lobe tips of pavement cells. Scale bars, 50 µm.

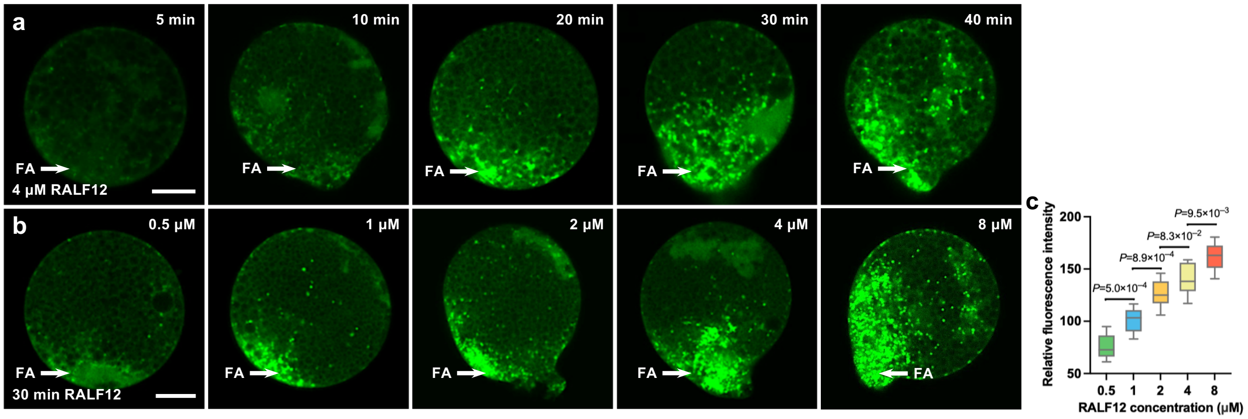


**Supplementary Fig. 5 RALF12 triggers high levels of granular ROS accumulation at the filiform apparatus region in a time- and concentration-dependent manner.** Synergid cells for RALF12 treatment were manually isolated from virgin ovules. **a** RALF12 treatment over time induces higher levels of granular ROS accumulation at the filiform apparatus area of synergid cells ultimately expanding within the whole cell. **b**,**c** Increasing concentrations of RALF12 significantly increases granular ROS accumulation. Data for relative ROS fluorescence intensity at the filiform apparatus region are present in box-and-whisker plots. Bottom and top of the box, 25th and 75th percentiles; centre line, 50th percentile; whiskers, minimum and maximum data. Statistical significance is determined by Student's *t*-test. Scale bars, 10 μm.

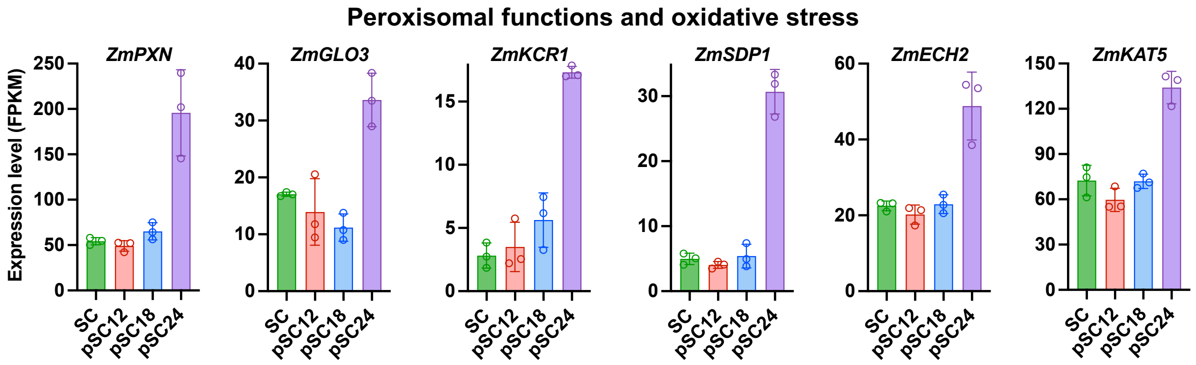


**Supplementary Fig. 6 Peroxisomal function-related genes are significantly activated indicating an oxidative stress status in synergids after 18 HAP.** Expression analysis of selected maize genes involved in peroxisomal functions including $NAD^+$ import (*ZmPXN*), 2-hydroxyacid oxidation (*ZmGLO3*), very long-chain fatty acids (VLCFAs) synthesis for peroxisomal metabolization (*ZmKCR1*) and fatty acid oxidation (*ZmSDP1*, *ZmECH2*, and *ZmKAT5*). Their expression pattern indicates an oxidative damage status in persistent synergid cells after 18 HAP.

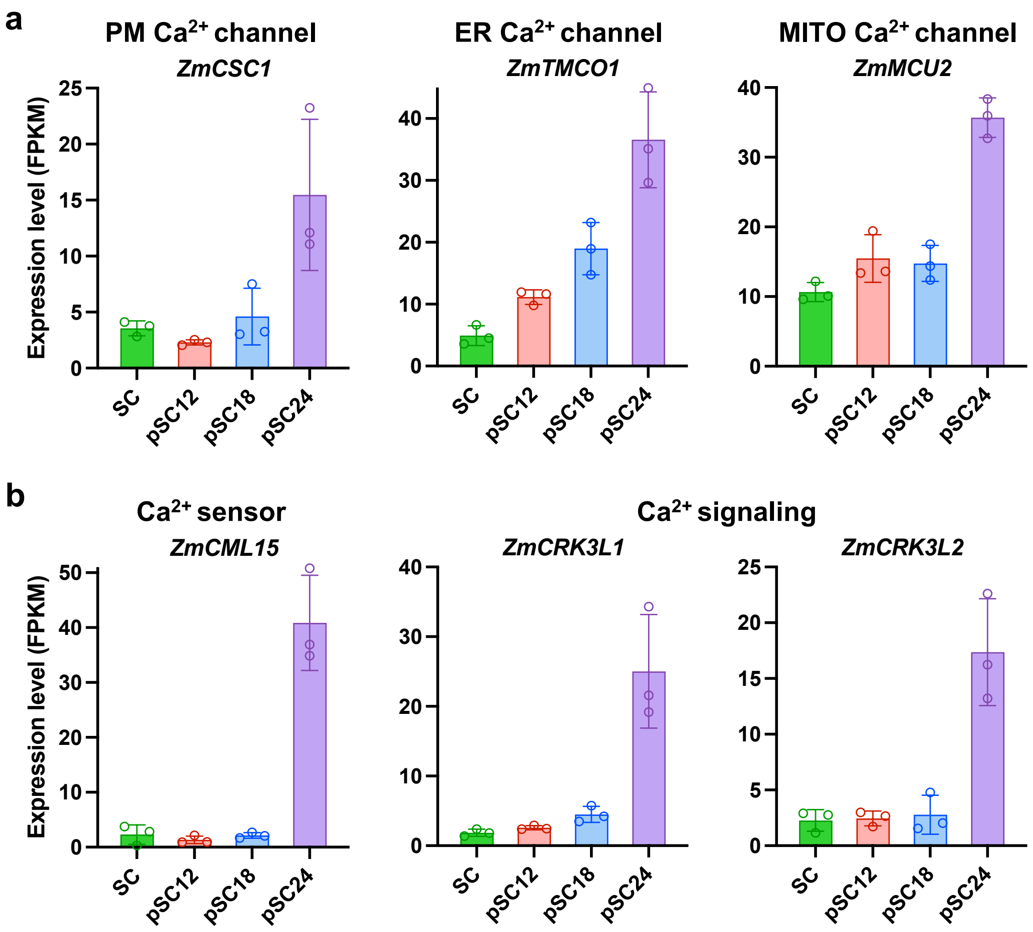


**Supplementary Fig. 7 Genes encoding $Ca^{2+}$ channels, $Ca^{2+}$ sensors, and $Ca^{2+}$ signaling components involved in senescence and cell death regulation are activated in synergids after 18 HAP. A** Activation of genes encoding a plasma membrane (PM) $Ca^{2+}$ permeable stress-gated cation channel (*ZmCSC1*), an ER membrane $Ca^{2+}$ load-activated $Ca^{2+}$ channel (*ZmTMCO1*), and a mitochondrial (MITO) inner membrane $Ca^{2+}$ uniporter (*ZmMCU2*) that mediates $Ca^{2+}$ uptake into mitochondria and activation of the cell death pathway. **b** Activation of genes encoding homologs of an *Arabidopsis* $Ca^{2+}$ sensor and a $Ca^{2+}$ signaling pathway kinases involved in senescence regulation.

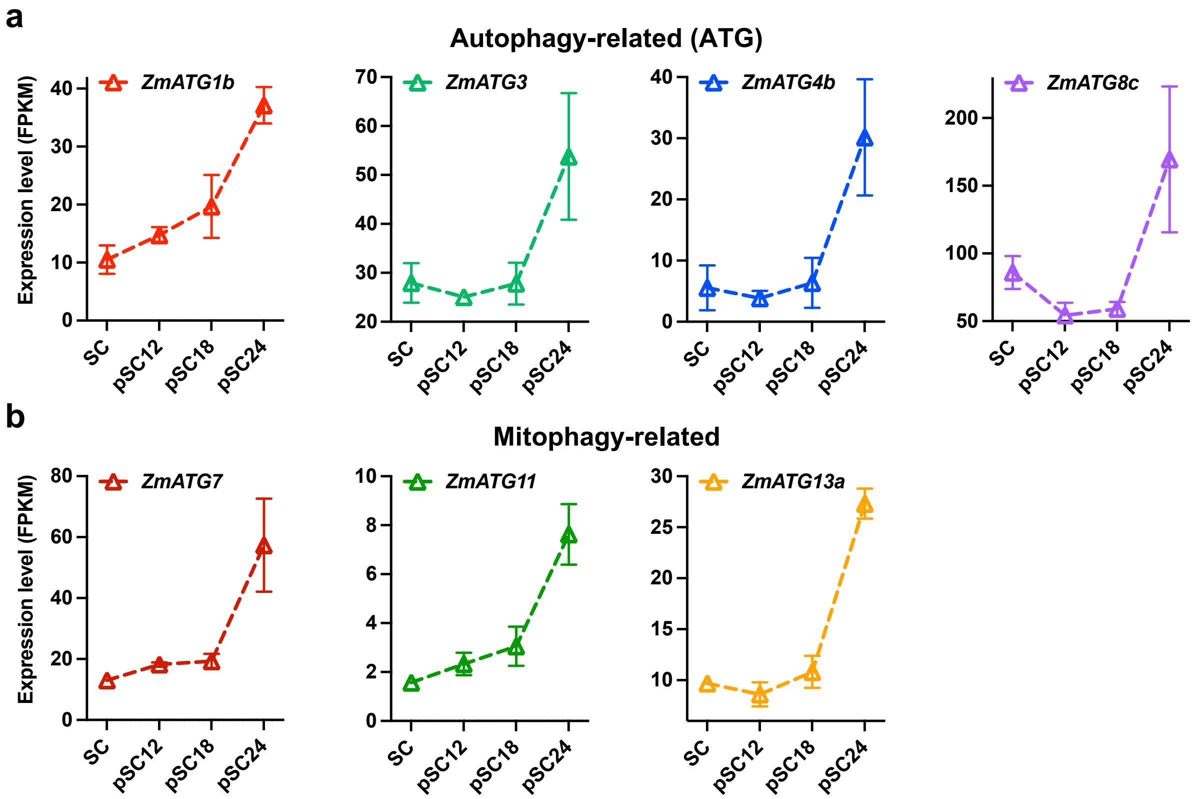


**Supplementary Fig. 8 Autophagy-related genes are activated with delay after fertilization in synergids for corpse clearance.** Indicated general autophagy-related genes (a) and mitophagy-related genes (b) are strongly activated at 18–24 HAP, shortly before persistent synergid cell elimination.

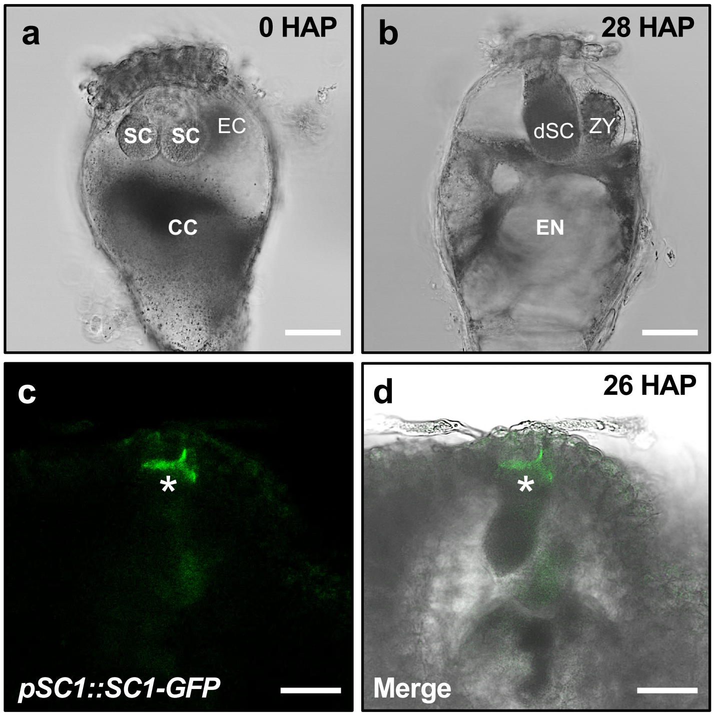


**Supplementary Fig. 9 Elimination of the persistent synergid cell through synergid-endosperm (SE) fusion at the end of PCD.** **a** Isolated maize embryo sac before pollination. **b** Isolated maize embryo sac at 28 HAP showing absence of the persistent synergid cell and moderate plasmolysis of a syncytium at the micropylar region. **c**,**d** An example of SC1-GFP signals at 26 HAP. Asterisk marks weak SC1-GFP signals at the residual filiform apparatus of the eliminated persistent synergid cell. Abbreviations: CC, central cell; dSC, degenerated synergid cell; EC, egg cell; EN, endosperm; SC, synergid cell; ZY, zygote. Scale bars, 50 µm.
